# Topoisomerases modulate the timing of meiotic DNA breakage and chromosome morphogenesis in *Saccharomyces cerevisiae*

**DOI:** 10.1101/672337

**Authors:** Jonna Heldrich, Xiaoji Sun, Luis A. Vale-Silva, Tovah E. Markowitz, Andreas Hochwagen

## Abstract

During meiotic prophase, concurrent transcription, recombination, and chromosome synapsis, place substantial topological strain on chromosomal DNA, but the role of topoisomerases in this context remains poorly defined. Here, we analyzed the roles topoisomerases I and II (Top1 and Top2) during meiotic prophase in *Saccharomyces cerevisiae*. We show that both topoisomerases accumulate primarily in promoter-containing intergenic regions of actively transcribing genes. Enrichment partially overlaps meiotic double-strand break (DSB) hotspots, but disruption of either topoisomerase has different effects during meiotic recombination. *TOP1* disruption delays DSB induction and shortens the window of DSB accumulation by an unknown mechanism. By contrast, temperature-sensitive *top2-1* mutants accumulate DSBs on synapsed chromosomes and exhibit a marked delay in meiotic chromosome remodeling. This defect results from a delay in recruiting the meiotic chromosome remodeler Pch2/TRIP13 but, unexpectedly, is not due to a loss of Top2 catalytic activity. Instead, mutant Top2-1 protein has reduced contact with chromatin but remains associated with meiotic chromosomes, and we provide evidence that this altered binding is responsible for the delay in chromosome remodeling. Our results imply independent roles for topoisomerases I and II in modulating meiotic recombination.

## Introduction

Topoisomerases preserve genome integrity by resolving topology-related strain and DNA entanglements associated with many cellular processes, including replication, transcription, and recombination (Pommier et al, 2016; Vos et al, 2011; Wang, 2002). To resolve strain, topoisomerases catalyze temporary breaks in the DNA. Type I topoisomerases make and re-ligate a single-strand break to relax strain in the DNA substrate, whereas type II enzymes catalyze DNA strand passage through a transient double-strand break (DSB). Topoisomerases are major chemotherapeutic targets that have been extensively studied in mitotically proliferating cells (Pommier et al, 2016). Comparatively less is known about the function of topoisomerases during meiosis.

Meiosis is a specialized type of cell division that is essential for sexual reproduction and allows for generation of genetic diversity. It involves a single round of DNA replication followed by two divisions that separate homologous chromosomes and sister chromatids, respectively, to produce four haploid cells from one diploid cell. In preparation for the first meiotic division, programmed DSB formation initiates exchange of DNA between homologous chromosomes by meiotic recombination (Borde & de Massy, 2013; Lam & Keeney, 2015a). This process allows for shuffling of genetic information and leads to the formation of crossovers, which help promote proper segregation of homologous chromosome pairs (Petronczki et al, 2003). Errors in this process can result in aneuploidy, infertility and congenital diseases, such as Down syndrome (Hassold & Sherman, 2000).

To support meiotic recombination, meiotic chromosomes assemble conserved loop-axis structures in which actively transcribing chromatin loops are anchored to a proteinaceous axial element (Sun et al, 2015; Zickler & Kleckner, 2015). As meiotic recombination progresses, axial elements of homologous chromosome pairs align and zip up to form a synaptonemal complex (SC) (Zickler & Kleckner, 2015). The SC limits further DSB induction and eases meiosis-specific repair restrictions to facilitate the repair of remaining DSBs before cells exit from meiotic prophase and initiate the first meiotic division (Subramanian et al, 2016; Thacker et al, 2014). Transcription, recombination and chromosome morphogenesis take place concurrently during meiotic prophase, but whether and how topoisomerases facilitate these processes remains poorly understood.

The meiotic functions of topoisomerases have been primarily elucidated in the context of the meiotic divisions where, similar to mitosis, topoisomerase II (topo II) has a major role in disentangling DNA to facilitate chromosome segregation (Gomez et al, 2014; Hartsuiker et al, 1998; Hughes & Hawley, 2014; Jaramillo-Lambert et al, 2016; Kallio & Lahdetie, 1996; Mengoli et al, 2014; Tateno & Kamiguchi, 2001).

All major topoisomerases are also present and active in meiotic prophase (Borde et al, 1999; Cobb et al, 1997; Stern & Hotta, 1983). The best-characterized is topoisomerase III, a type I topoisomerase, which decatenates recombination intermediates during meiotic prophase (De Muyt et al, 2012; Gangloff et al, 1999; Kaur et al, 2015; Tang et al, 2015). Only minor meiotic defects have been reported upon inactivation of topo I, including increased gene conversion events in the ribosomal DNA of *Saccharomyces cerevisiae* (Christman et al, 1988) and mild defects in chromosome pairing in mice (Cobb et al, 1997; Handel et al, 1995). Somewhat more is known about topo II, which localizes diffusely to prophase chromatin in a variety of organisms (Cheng et al, 2006; Cobb et al, 1999; Jaramillo-Lambert et al, 2016; Zhang et al, 2014), with enrichment along chromosome axes noted in some cases (Klein et al, 1992; Moens & Earnshaw, 1989). In *Saccharomyces cerevisiae*, topo II cleaves preferentially in promoter regions during meiotic prophase (Borde et al, 1999) and contributes to proper spacing of crossover events (Zhang et al, 2014). Aberrant recombination upon chemical inhibition of topo II has also been noted in mouse spermatocytes (Russell et al, 2000). In addition, topo II helps resolve chromosome interlocks in *Arabidopsis* (Martinez-Garcia et al, 2018). Possibly related to this function, *Saccharomyces cerevisiae* topo II mutants arrest at the end of meiotic prophase in a DSB-dependent manner despite the appearance of mature recombinants (Rose & Holm, 1993; Rose et al, 1990; Zhang et al, 2014). However, an in-depth analysis of topoisomerase I and II distribution on prophase chromosomes has so far not been performed.

In this study, we used chromatin immunoprecipitation and deep sequencing (ChIP-seq) to determine the meiotic distribution of topoisomerases I and II (encoded by *TOP1* and *TOP2*) in *S. cerevisiae*. We show that both topoisomerases are primarily enriched in promoter-containing intergenic regions (IGRs) and that enrichment increases upon meiotic entry and correlates with transcriptional activity. Despite the comparable binding patterns, *top1* and *top2* mutations have different effects on meiotic prophase. Deletion of *TOP1* alters the timing of meiotic DSB formation, whereas *top2-1* mutants primarily show defects in chromosome morphogenesis.

## Results

### Topoisomerases are enriched in promoter-containing IGRs

We examined the chromosomal association of yeast Top1 and Top2 in a synchronous meiotic time course. Immunofluorescence analysis of chromosome spreads showed that both proteins form foci on chromatin that are detectable at all stages of meiotic prophase as well as prior to meiotic induction (**Figures S1a-b**). As chromosomes compact to form the SC, Top1-13myc and Top2 are detectable both on chromatin loops and in the vicinity of the chromosome cores, as marked by the SC protein Zip1. Both proteins, especially Top1-13myc, are also present in the nucleolus, which is devoid of Zip1 staining (arrowhead, **Figure S1a**).

To obtain more detailed spatial information, we analyzed the genomic distribution of Top1 and Top2 by ChIP-seq at the time of maximal meiotic DSB formation (3h after meiotic induction at 30°C (Falk et al, 2010)). This analysis revealed that Top1 and Top2 bind in a similar pattern (**Figure 1a** and **S1c**, correlation score = 0.76). Metagene analysis showed a particular enrichment in the promoter regions upstream of gene bodies (**Figure 1b**), consistent with analyses of Top2 in vegetative cells (Bermejo et al, 2007; Gittens et al, 2019; Sperling et al, 2011) and the analysis of Top2 cleavage complexes in meiosis (Borde et al, 1999). The enrichment downstream of open reading frames (ORFs) is primarily a consequence of the promoter of the next gene. When signals were parsed into divergent, tandem, and convergent intergenic regions (IGRs), topoisomerase enrichment was observed in IGRs containing at least one promoter, with the strongest signal observed for divergent IGRs (**Figures 1c** and **S1d**). By contrast, convergent IGRs showed only weak signal. The dip between convergent gene pairs resembles the binding of axial-element proteins (Sun et al, 2015), suggesting a possible influence of the axis on topoisomerase binding in these regions. In promoter-containing IGRs, both topoisomerases were broadly bound in the intergenic space delimited by the two flanking genes (**Figure 1d-e**), although topoisomerase enrichment appeared comparatively reduced in narrow IGRs (**Figure 1f-g**). Thus, the size of promoter-containing IGRs may influence topoisomerase recruitment.

**Figure 1.**
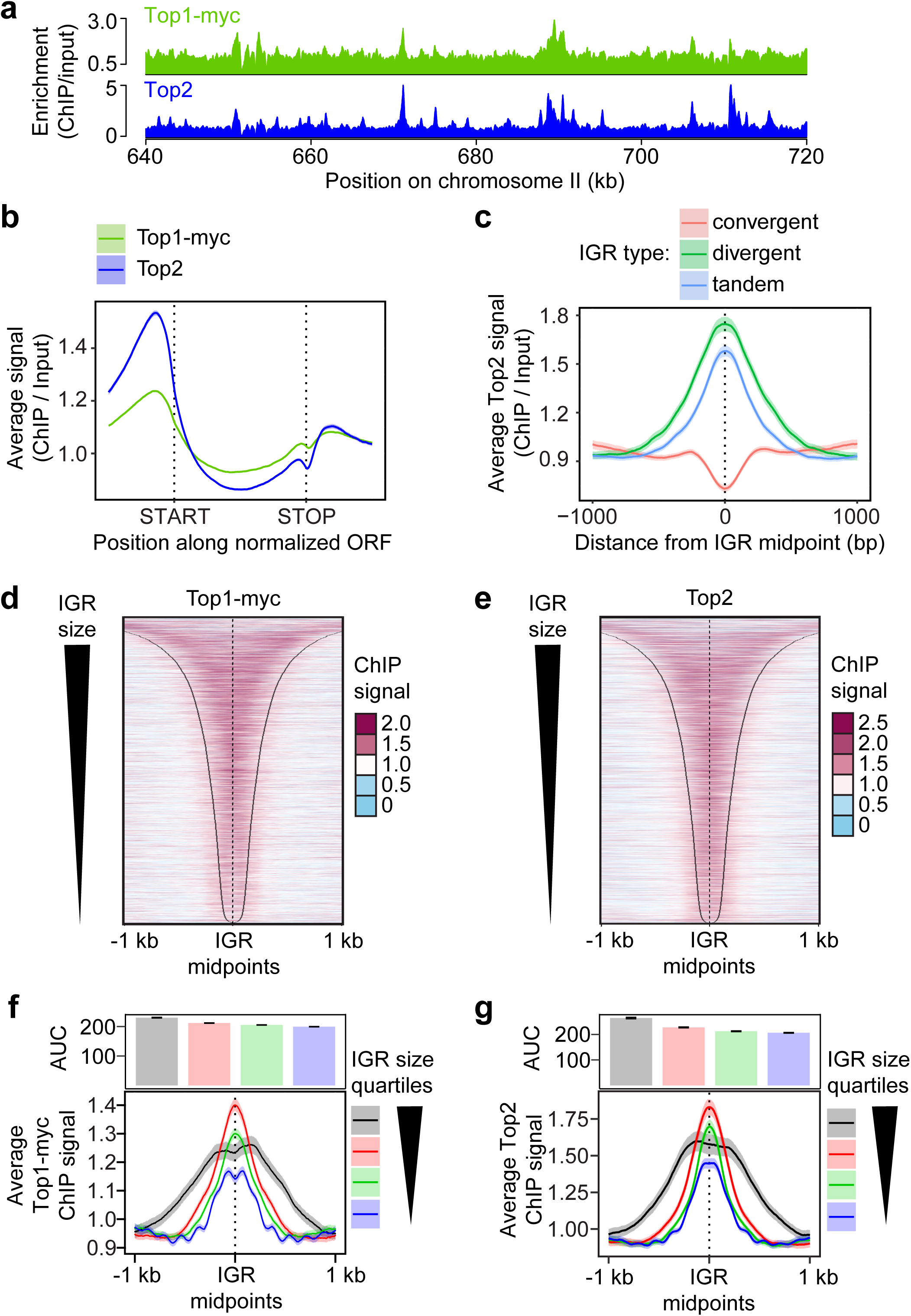
Topoisomerases are enriched in promoter-containing IGRs. **(a)** Wild-type genomic distribution of Top1-13myc and Top2 on a section chromosome II at the time of meiotic DSB formation (3h after meiotic induction) as measured by ChIP-seq. **(b)** Metagene analysis Top1 and Top2 enrichment. Start and end positions of the scaled ORFs are indicated and the flanking regions represent the space up- and downstream of a given ORF equal to half the width of the ORF. **(c)** Top2 enrichment centered at the midpoints of IGRs parsed into divergent, tandem, and convergent regions. The 95% confidence interval is shown for average lines in panels (b) and (c). Heat maps of **(d)** Top1 and **(e)** Top2 localization centered at midpoints of all promoter-containing IGRs (i.e. divergent and tandem), sorted by size of the IGR. Black lines delineate start or end of ORFs bordering the IGR. Average **(f)** Top1 and **(g)** Top2 signal in quartiles based on IGR size. Quartile ranges are <294 bp (blue), 294 - 453 bp (green), 454 - 752 bp (red), and >752 bp (black). Signals are centered at IGR midpoints and extended 1kb in each direction. The 95% confidence interval for the average lines is shown for panels (f) and (g). The area under the curve (AUC) is quantified in the bar plots above the quartile signal plots. Black bars represent standard error for each quartile. All quartiles are significantly different (*P* < 0.0001, Mann-Whitney-Wilcoxon test with Bonferroni correction).

### Topoisomerase binding correlates with gene expression

We tested whether meiotic topoisomerase enrichment in promoter-containing IGRs is linked to the expression of the flanking genes by performing mRNA-seq analysis 3h after meiotic induction. This analysis revealed a weak positive correlation between topoisomerase binding and steady-state mRNA levels (**Figure 2a** and **S2a-b,** correlation score ∼ 0.2 for both topoisomerases). The correlation was strongest in promoter regions but extended across ORFs for the most highly expressed quartile, especially for Top1-myc, consistent with increased buildup of topological stress on highly expressed genes (Teves & Henikoff, 2014).

**Figure 2.**
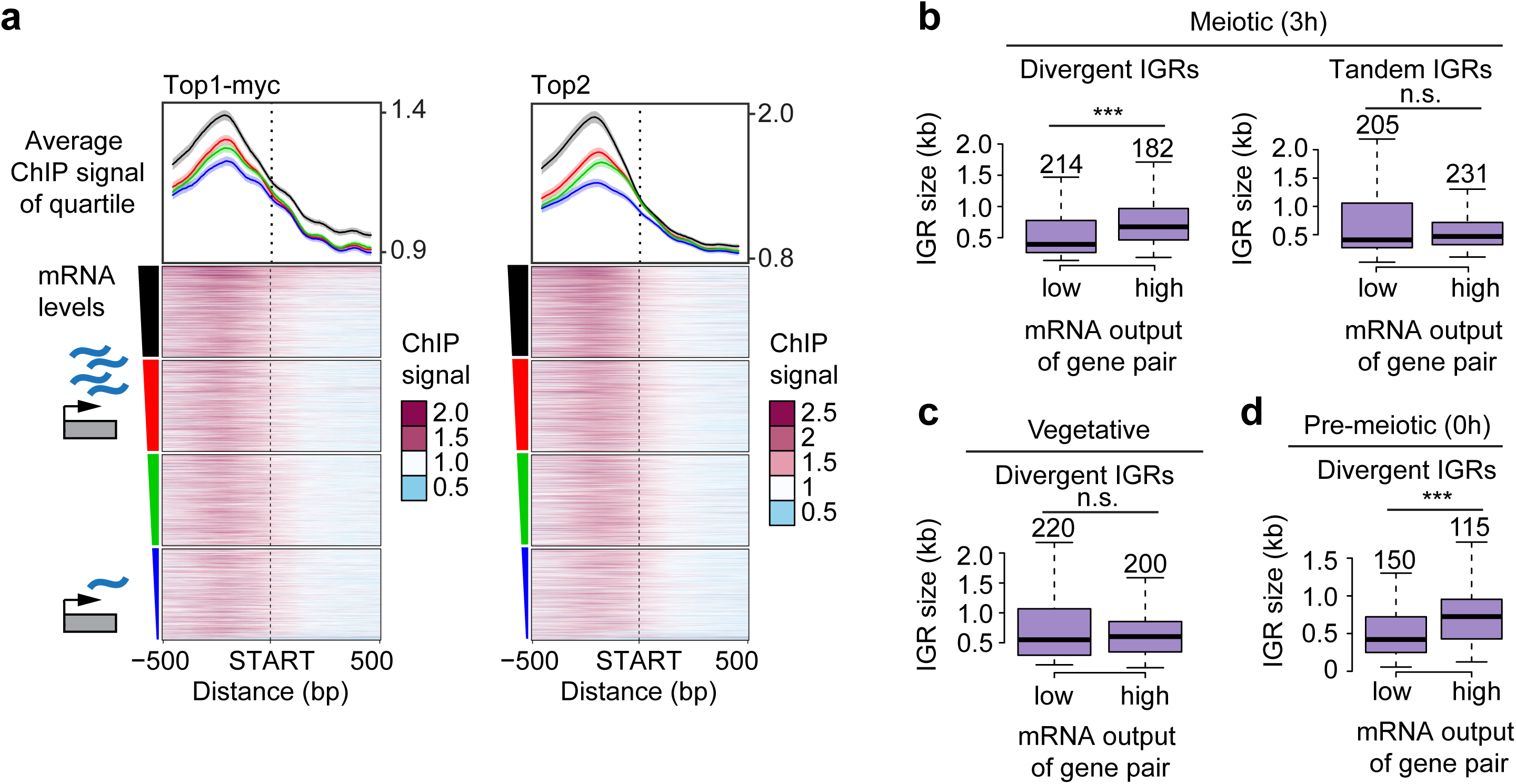
Topoisomerase enrichment correlates with mRNA levels. **(a)** Top1 and Top2 localization 500 bp up- and downstream of starts of ORFs, sorted based on the amount of steady-state mRNA of the associated gene. The average of each quartile is plotted above the heat maps. The colors of the lines correspond to the color segments beside the 4 quartiles of transcriptional activity. The 95% confidence interval for the average quartile lines is shown. **(b)** Box plots showing the size distribution of divergent and tandem IGRs during meiosis for gene pairs with either extremely high or low levels of associated mRNA. The size of each group was chosen to achieve similar numbers of gene pairs, which are noted above the respective box, and vary due to the changes in the transcriptional program as well as the data set used. Significance was determined by unpaired, two-sided Wilcoxon test. **(c-d)** Similar analysis as in (b) for divergent IGRs in vegetative (c) and pre-meiotic (d) cells using data from (Cheng et al, 2018). Note that gene pair identity changes as a function of the transcriptional program in the different developmental stages. \*\*\**P* < 0.001 and n.s. P-value=0.71 (b) and P-value=0.54 (c), Mann-Whitney-Wilcoxon test.

Unexpectedly, gene expression levels during meiosis are also positively correlated with the size of divergent IGRs. Comparing divergent IGR sizes as a function of mRNA levels showed that the median IGR size of gene pairs in which both genes are among the most highly expressed is nearly twice as large as the median IGR size of gene pairs in which both genes are among the most lowly expressed (674 bp compared to 393 bp) (**Figure 2b**). This bias likely accounts for the wider topoisomerase profile in the most highly expressed quartile (**Figure 2a**). Tandem IGRs did not show this bias (**Figure 2b**), indicating that this feature is linked to relative gene arrangement. We confirmed this association by analyzing published mRNA-seq time course data (Cheng et al, 2018). This analysis showed that highly expressed gene pairs are preferentially associated with large divergent IGRs throughout meiosis but not in vegetative cells (**Figures 2c** and **S2c**). The effect is already seen prior to meiotic entry (T=0h) and in non-meiotic *MATa/a* cells in sporulation medium (**Figures 2d** and **S2c**), suggesting that it is linked to the starvation regime used to induce synchronous meiosis in yeast. This link between IGR size and expression levels may also contribute to the apparently lower enrichment of topoisomerases in narrow IGRs (**Figure 1f-g**).

### Meiotic entry leads to a buildup of Top2 on meiotic chromosomes

To test if meiotic chromosome morphogenesis impacts topoisomerase recruitment, we followed Top2 enrichment as cells transition from pre-meiotic G1 (T=0h) into meiotic prophase (T=3h) using spike-in normalized ChIP-seq analysis (SNP-ChIP (Vale-Silva et al, 2019)). This analysis showed an overall ∼35% increase in Top2 binding across the genome as cells transition into meiotic prophase (**Figure 3a**). This increase is not a sign of ongoing replication, because under our experimental conditions pre-meiotic S phase is largely complete before the 3-hour time point (e.g. **Figure 4a**). In addition, the increase is not linked to axis morphogenesis because it also occurred in mutants lacking the cohesin Rec8 (**Figure 3a**), which is required for axial-element assembly (Klein et al, 1999). Interestingly, chromatin-associated Top2 levels increased a further ∼30% when wild-type cells were entering prophase at elevated temperature (34°C). This additional accumulation raises the possibility that elevated temperature increases the topological strain of meiotic chromosomes.

**Figure 3.**
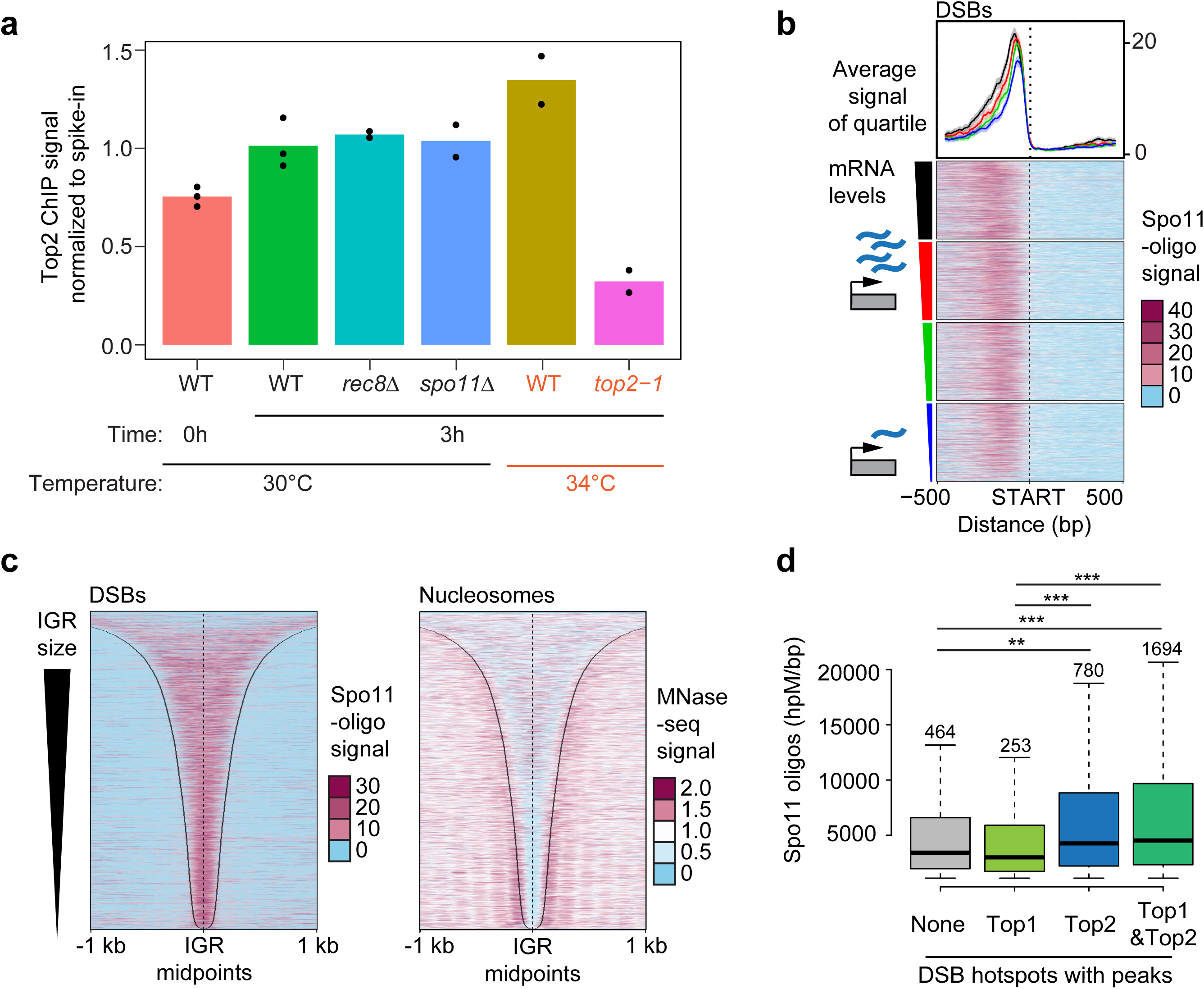
Sites of topoisomerase enrichment partially overlap with DSB hotspots. **(a)** Comparison of the level of total chromosomal association of Top2 on pre-meiotic (0h) and meiotic chromosomes (3h after meiotic induction) as determined by SNP-ChIP spike-in analysis. Top2 levels are determined for meiotic chromosomes for various mutant backgrounds (*rec8Δ* and *spo11Δ*) as well as at 34°C (wild type and *top2-1*). Points represent individual replicate values and bars represent average. Values for each experimental replicate are normalized to the average wild-type meiotic (3h) levels. **(b)** Heat maps of Spo11-oligo signal 500 bp up- and downstream of starts of ORFs, sorted based on the amount of steady-state mRNA of the associated gene. Colored triangle segments indicate 4 quartiles of transcriptional activity. The average and 95% confidence interval of each quartile is plotted above the heat maps. Heat maps of **(c)** Spo11-oligo signal and nucleosome signal determined by MNase-seq across all promoter regions sorted by IGR size. Black lines delineate IGR borders. **(d)** Comparison of hotspot activity based on whether hotspots overlap with a significant peak of either no topoisomerase, Top1, Top2, or both Top1 and Top2. Significant peaks were determined by MACS (see Methods). Number of hotspots in each group is labeled above the respective box in the plot. \*\*\**P* < 0.0001, \*\**P* < 0. 01, Mann-Whitney-Wilcoxon test with Bonferroni correction.

**Figure 4.**
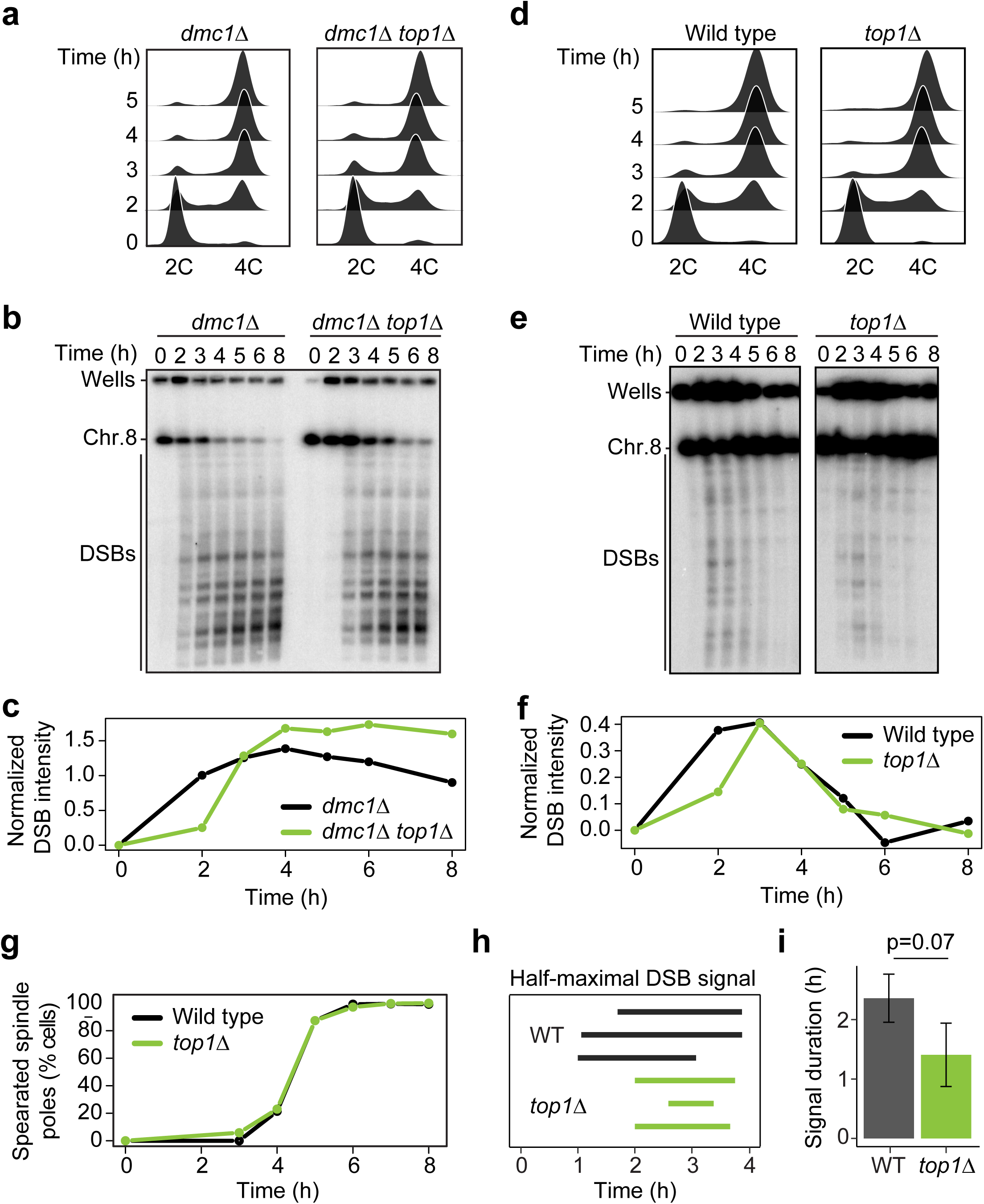
Loss of *TOP1* shortens the interval of DSB activity. **(a)** DNA content of *dmc1Δ* and *dmc1Δ top1Δ* cells as determined by flow cytometry. Samples were taken at the indicated time points. **(b)** PFGE/Southern analysis of DSBs along chromosome VIII in *dmc1Δ* and *dmc1Δ top1Δ* cells. Results were consistent across three experiments. **(c)** Quantification of DSB signal in (b) calculated as fraction of total signal (parental and DSB) after background subtraction. **(d-f)** Time course analysis of wild-type and *top1Δ* cells. The analyses are the same as in (a-c) (n=3). **(g)** Spindle pole separation as measured by anti-tubulin immunofluorescence of wild-type (black) and *top1Δ* (green) cells (n=200). **(h-i)** The time after meiotic induction (h) and the duration of the time interval (i) in which the DSB signal was above half the maximum value for wild type and *top1Δ* in three experiments. Error bars in (i) are standard deviation. P=0.07, unpaired t-test.

### Topoisomerase enrichment is correlated with meiotic DSB hotspot activity

As promoter-containing IGRs frequently serve as meiotic DSB hotspots (Lam & Keeney, 2015b; Pan et al, 2011), we compared the genomic distribution of Top2 with the sites of activity of Spo11, the topoisomerase II-related protein that catalyzes meiotic DSB formation (Bergerat et al, 1997; Keeney et al, 1997). For this analysis, we used available high-resolution sequencing data of Spo11-associated oligonucleotides (Spo11-oligos), which report on Spo11 cleavage activity (Pan et al, 2011; Thacker et al, 2014). Consistent with previous work (Blitzblau et al, 2007; Gittens et al, 2019), Spo11-oligo signal is strongest in the promoter regions of highly expressed genes (**Figure 3b**). However, unlike for topoisomerase enrichment, there is no consistent correlation between Spo11 and gene expression (**Figure S3b**, correlation score = 0.05) (Zhu & Keeney, 2015). Rather, the correlation plot suggests that only the promoter regions of the most highly expressed genes have a greater likelihood of being strong DSB hotspots and drive the positive slope.

Other aspects Spo11 cleavage patterns also differed from the patterns observed for topoisomerase association. Most notably, Spo11 cleavage activity is more focused than topoisomerase binding signal (compare **Figures 1d,e** and **3c**) (Pan et al, 2011) and shows a different correlation with IGR size (**Figure 3c, S3d**). Some of these differences may be due to assay differences, as Spo11-oligos map precise cleavage sites, whereas ChIP-seq analysis maps broader regions of association based on formaldehyde crosslinking and DNA fragmention. Nonetheless, these observations argue against a single shared mechanism driving topoisomerase recruitment and Spo11 activity. Accordingly, SNP-ChIP analysis showed that Top2 binding levels are unchanged in *spo11Δ* mutants (**Figure 3a**), indicating that the prophase enrichment of Top2 on meiotic chromosomes occurs upstream or in parallel to Spo11 activity.

It is possible that topoisomerases and Spo11 respond similarly to the local chromatin environment at promoter-containing IGRs. Indeed, when we analyzed relative topoisomerase enrichment after splitting DSB hotspots into quintiles based on Spo11 activity, we observed a weak but significant correlation between Top1 and Top2 enrichment and Spo11 activity based on 95% confidence intervals (**Figure S3c**). To further probe this link, we conducted the inverse analysis. We compared the average Spo11 activity of hotspots overlapping with a strong peak of Top1 or Top2 with those that did not (**Figure 3e**). This analysis revealed that DSB hotspot activity is elevated at hotspots that exhibit significant Top2 enrichment. By contrast, Top1 enrichment showed no correlation regardless of Top2 enrichment, suggesting that Top1 and Top2 interact differently with DSB hotspots.

### Loss of *TOP1* shortens the interval of meiotic DSB formation

To test if topoisomerase enrichment at meiotic DSB hotspots has functional consequences for meiotic recombination, we monitored meiotic DSB activity in cells in which either topoisomerase had been inactivated. After collecting DNA from a synchronous meiotic time course, we followed DSB formation by pulsed-field gel electrophoresis (PFGE) and Southern blotting for chromosome VIII. To analyze accumulation of DSBs, we used mutants with a *dmc1Δ* background, which are unable to repair DSBs (Bishop et al, 1992).

Flow cytometry analysis showed that *top1Δ dmc1Δ* mutants underwent pre-meiotic DNA replication with wild-type kinetics (**Figure 4a**). However, the appearance of meiotic DSB bands was delayed by approximately 1 hour (**Figure 4b-c**). This delay was reproducible and does not reflect a reduced ability to form DSBs, because *top1Δ dmc1Δ* mutants ultimately reached and even exceeded the DSB levels of *dmc1Δ* mutants. It is also not a consequence of the *dmc1Δ* background because repair-competent (*DMC1*) *top1Δ* mutant cells showed a similar delay in DSB induction despite apparently normal replication timing (**Figure 4d-f**). Intriguingly, despite the delay in initiation, the DSB signal of *top1Δ* mutants disappeared with near wild-type kinetics, and spindle formation occurred at the same rate as wild type (**Figure 4e-g**). Quantifying the duration of half maximal breakage showed that the window of DSB formation and repair is noticeably reduced in the absence of Top1, although the difference did not reach the p<0.05 cutoff (t-test, p=0.07; **Figure 4h-i**). As *top1Δ* mutants are fully competent to form DSBs (**Figure 4b-c**), the shorter interval of DSB formation likely either reflects a shortened window of opportunity for DSB formation in the absence of Top1 or accelerated DSB turnover. Notably, SC morphology and spore viability of *top1Δ* mutants were indistinguishable from wild type (**Figure S4a-c**), indicating that despite the shortened DSB interval, sufficient crossovers formed to support faithful meiotic chromosome disjunction.

### *TOP2* inactivation causes persistent DSB signal

As *TOP2* is an essential gene, we used the thermo-sensitive *top2-1* mutant, which exhibits no detectable catalytic activity above 30°C (DiNardo et al, 1984). For experiments utilizing this mutant, wild-type and mutant cells were induced to enter meiosis at room temperature and then shifted to 34°C one hour after meiotic induction. Flow cytometry analysis of *top2-1* showed less synchronous entry into meiosis compared to control cells, likely because of the slower growth of *top2-1* mutants (**Figure 5a,d**). DSB formation nevertheless initiated on time or even slightly earlier than in control cells, regardless of whether cells were competent for repair, and reached levels similar to wild type in a *dmc1Δ* mutant background (**Figures 5b-c** and **S5a-b**). Moreover, Spo11-oligo levels were comparable to wild type (**Figure S5c**). No breaks were observed in *top2-1* mutants lacking *SPO11* (**Figure S5d**), indicating that DSB signal is not the result of breaks arising from defects in DNA replication.

**Figure 5.**
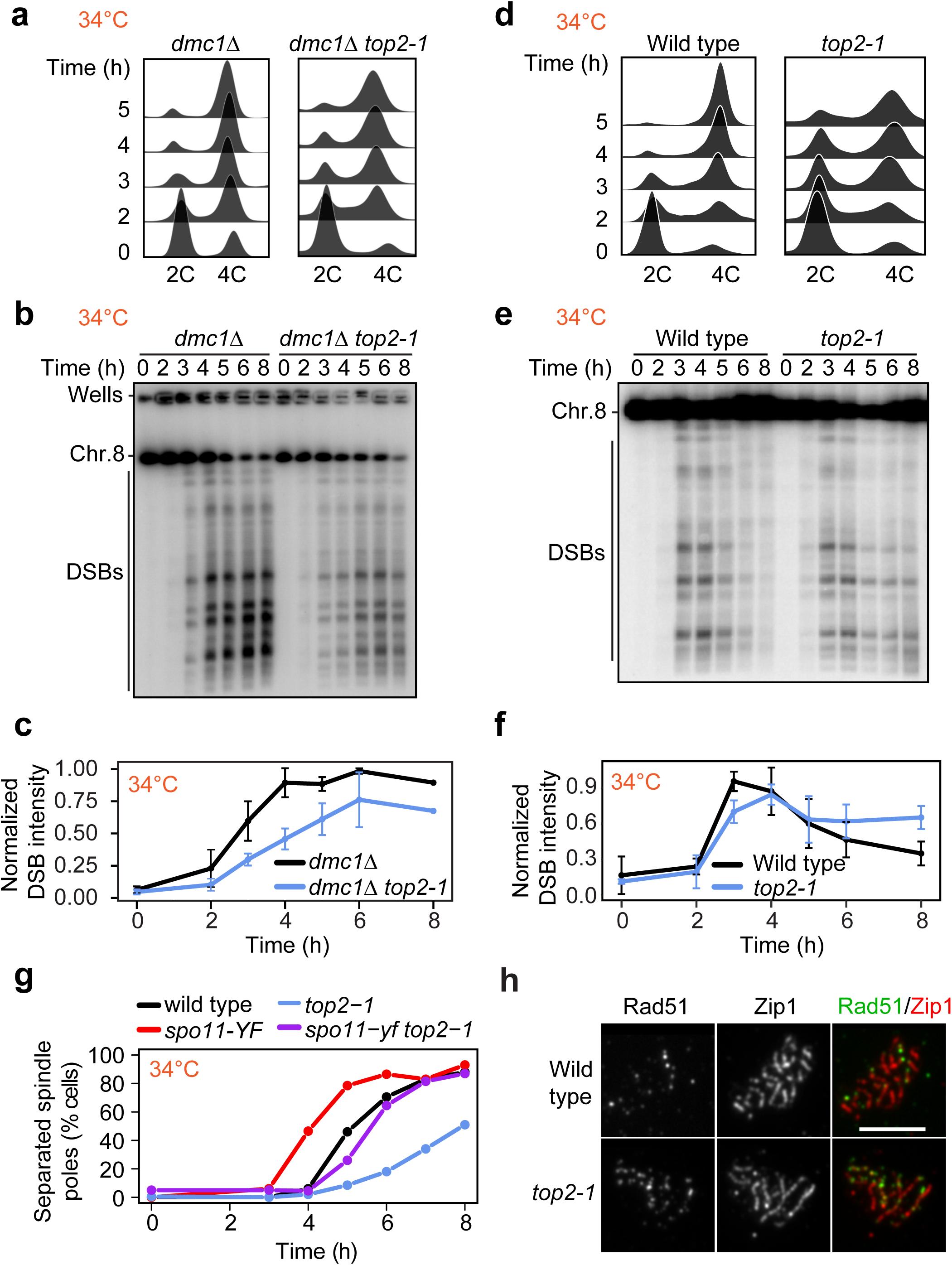
DSBs persist late into prophase in *top2-1* mutants. **(a)** DNA content of *dmc1Δ* and *dmc1Δ top2-1* cells as determined by flow cytometry. Samples were taken at the indicated time point. **(b)** Southern blot analysis of DSBs throughout a meiotic time course across chromosome VIII in wild-type and *top2-1* cells. Cultures were shifted to the restrictive temperature (34°C) 1h after meiotic induction. **(c)** Background signal was subtracted from DSB signal and each experiment was normalized to the max signal of the control. Average and standard deviation for three experiments are plotted. **(d-f)** Time course analysis of wild-type and *top2-1* cells. The analysis methods are the same as in (a-c). **(f)** Spindle pole separation as measured by anti-tubulin immunofluorescence of wild-type (black), *top2-1* (blue), *spo11-Y135F* (red), and *top2-1 spo11-Y135F* (purple) cells (n=200 for each data point). **(h)** Representative images of immunofluorescence staining for Rad51 (green) and Zip1 (red) on chromosome spreads of wild type and the *top2-1* mutant at the restrictive temperature (34°C) 4-5 hr after meiotic induction. Scale bar is 5 μm.

In repair-competent *top2-1* mutants (*DMC1*), DSB signal remained detectable after wild-type cells had completed DSB repair (**Figure 5e-f**), leading to a delayed prophase exit as assayed by spindle formation (**Figure 5g**). To determine to what extent the persistent DSB signal is a consequence of the poorer synchrony of the *top2-1* mutant, we analyzed the kinetics of spindle-pole separation in the presence of a catalytically inactive *spo11-Y135F* mutation, which prevents DSB formation (Keeney et al, 1997). As expected, the *spo11-Y135F* mutant completed meiosis I more rapidly than wild type because prophase is shortened in this mutant (Kee & Keeney, 2002). In a *top2-1 spo11-Y135F* double mutant, spindle poles separated approximately 1.5 hours slower than in the *spo11-Y135F* single mutant (**Figure 5g**). These data indicate that poorer synchrony accounts for 1.5 hours of the prophase delay of the *top2-1* mutant. However, DSB signals persisted unchanged for at least 3 hours (**Figure 5e-f**), suggesting that part of the persistent DSB signal is the result of slower DSB turnover. In line with this interpretation, we observed an elevated number of Rad51 foci on synapsed chromosomes, indicating the presence of DSBs late into prophase (**Figure 5h**; see also **Figure 7c**).

### Inactive Top2 protein shows reduced and altered binding to chromatin

To better understand the effects of the *top2-1* mutant, we analyzed the chromosomal association of mutant Top2 protein during meiosis. Immunofluorescence staining for Top2 on chromosome spreads showed foci localizing abundantly to meiotic chromosomes in the *top2-1* mutant (**Figure 6a**), demonstrating that despite the loss of catalytic activity above 30°C (DiNardo et al, 1984), mutant Top2 protein retains the capacity to bind to meiotic chromosomes. However, SNP-ChIP analysis showed a more than 4-fold drop in signal (**Figure 4a**), indicating that the amount of mutant Top2 that can be crosslinked to DNA is strongly reduced. ChIP-seq analysis in *top2-1* mutants revealed a loss of Top2 binding from most promoter-containing IGRs (**Figure 6b-c** and **S6a-b**), whereas binding appeared less affected at sites enriched for the meiotic chromosome axis factor Red1 (arrow heads, **Figure 6b**). Accordingly, we observed a relative enrichment of mutant Top2 in convergent regions (**Figure 6c**), which are preferred sites of Red1 binding (Sun et al, 2015). This spatial association may reflect interactions of mutant Top2 protein with the meiotic chromosome axis.

**Figure 6.**
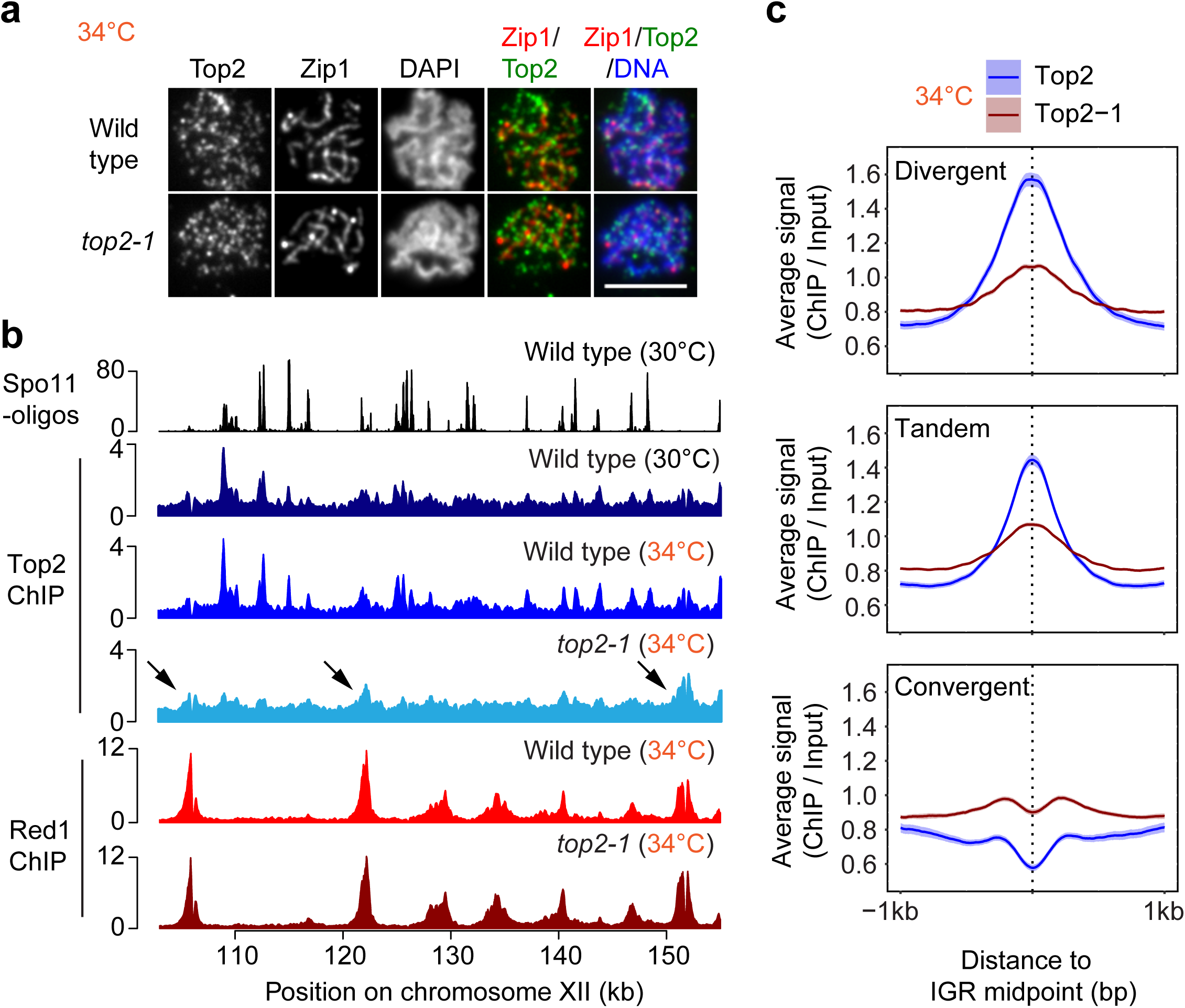
Top2-1 binding is retained at sites of Red1 enrichment. **(a)** Immunofluorescence staining for Top2, Zip1, and DAPI on chromosome spreads of wild type and the *top2-1* mutant at the restrictive temperature (34°C) 4-5 hr after meiotic induction. Scale bars are 5 μm. **(b)** ChIP-seq analysis of Top2 and the axis protein Red1 at 34°C in wild type and *top2-1* mutants shown for a representative region on chromosome XII and compared to Spo11 cleavage patterns at 30°C (Thacker et al, 2014) and Top2 binding at 30°C. Arrows mark Top2 peaks that remain in the *top2-1* mutant. These peaks show overlap with strong Red1 peaks. **(c)** Average Top2 binding in wild type and the *top2-1* mutant at the restrictive temperature at IGRs based on orientation of genes. The 95% confidence interval for the average lines is shown.

### Mutant Top2 interferes with synapsis-associated chromosome remodeling

Given the important role of chromosome structure in guiding meiotic DSB repair (Zickler & Kleckner, 2015), we asked whether the DSBs on synapsed chromosomes in *top2-1* mutants (**Figure 5h**) could be related to defects in chromosome morphogenesis. ChIP-seq analysis of Red1 indicated that the distribution of axis attachment sites was unaltered in *top2-1* mutant cultures (**Figure 6b**). To assay meiotic chromosome structure at the single-cell level, we stained chromosome spreads for the structural components Hop1 and Zip1. Hop1 is recruited to chromosomes prior to DSB formation and is removed at the time of repair, as the SC component Zip1 is deposited onto the chromosomes (Smith & Roeder, 1997; Subramanian et al, 2016). As a result, Hop1 and Zip1 show an alternating pattern on wild-type chromosomes (**Figure 7a**) (Börner et al, 2008). By contrast, Hop1 and Zip1 signals exhibited substantial overlap on *top2-1* chromosomes (**Figure 7a**). Furthermore, DAPI staining of *top2-1* mutants revealed the accumulation of chromosomes with characteristic parallel “train tracks”, which rarely appear in wild-type nuclei (**Figure 7b**).

**Figure 7.**
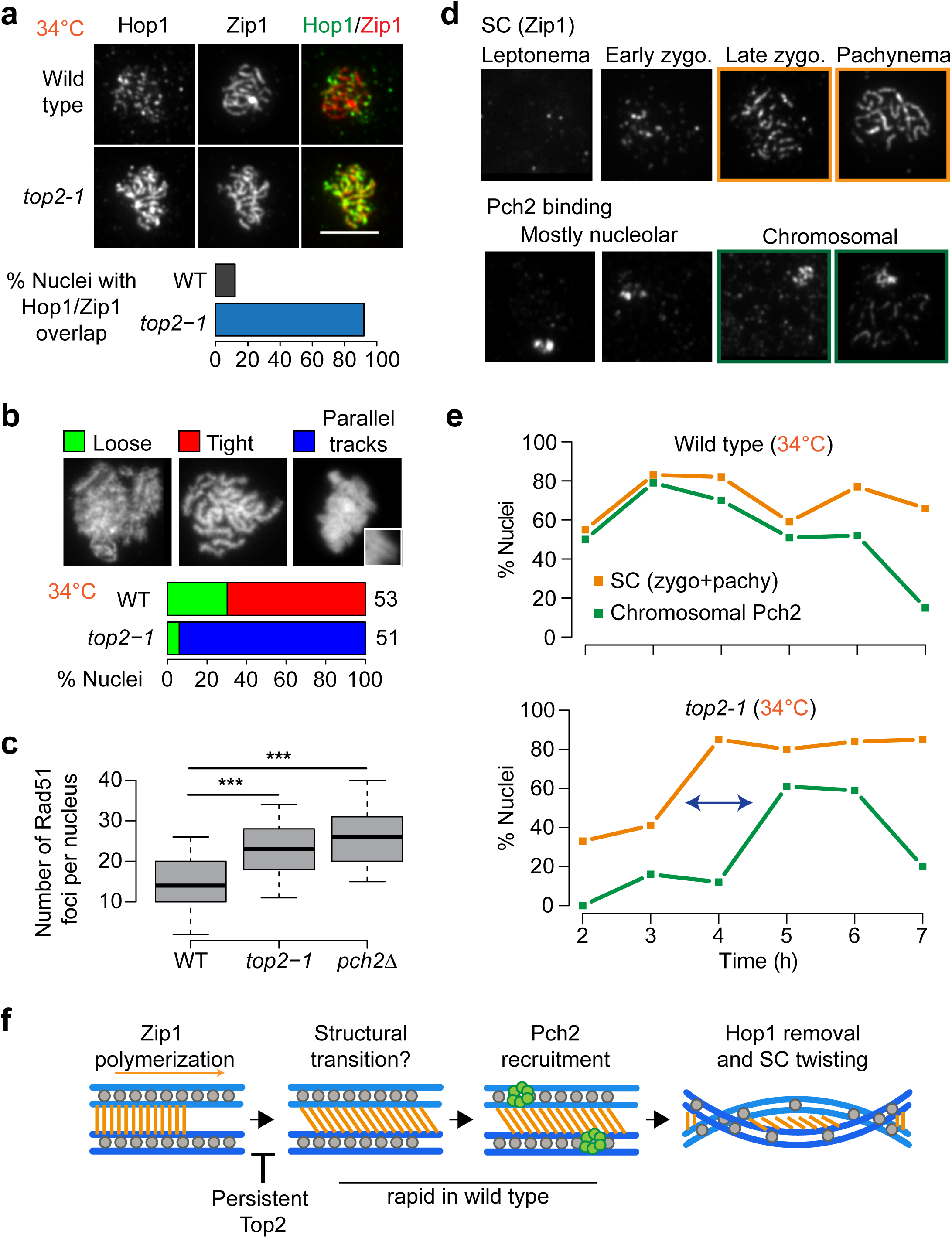
Mutant Top2 delays meiotic chromosome remodeling. **(a)** Immunofluorescence staining for Hop1 and Zip1 on chromosome spreads of wild type and the *top2-1* mutant during meiosis at 34°C. Scale bars are 5 μm. Bar graph shows the percent of nuclei with extensive Hop1 and Zip1 overlap in wild-type and *top2-1* cells. **(b)** Quantification of chromosomal phenotypes as determined by DAPI staining of chromosome spreads containing tracks of Zip1 marking late prophase. Images show representative examples of chromosomal phenotypes. Inset shows magnification of a typical train-track conformation. Analysis is shown for wild type and *top2-1*. The number of nuclei counted is indicated next to the respective bar in the plot. **(c)** Quantification of Rad51 foci on fully synapsed chromosomes (as determined by Zip1 staining on chromosome spreads) in wild type and *top2-1* and *pch2Δ* mutants at 34°C. \*\*\**P* < 0.0001, n.s. P-value=0.6, Mann-Whitney-Wilcoxon test with Bonferroni correction. **(d-e)** Quantification of immunofluorescence staining for Pch2 and Zip1 on wild-type and *top2-1* chromosome spreads throughout a meiotic time course at 34°C. **(d)** Representative images of Zip1 staging and Pch2 binding. **(e)** Orange lines show % cells with late zygotene or pachytene morphology at indicated time points. Green lines show % cells with abundant Pch2 staining on chromosomes in addition to the strong nucleolar signal. **(f)** Model for structural dynamics of the SC. Following Zip1 polymerization, a structural transition leads to the recruitment of Pch2, perhaps by making chromosomal Hop1 accessible as a substrate for Pch2. Following Hop1 removal, the chromosomes lose their train-track morphology, likely by twisting, as seen in many organisms. Aberrant Top2 binding inhibits the structural transition, leading to delayed Pch2 recruitment, extended Hop1 binding, and extended train-track morphology of the aligned chromosomes.

We observed similar, albeit weaker, phenotypes when strains carrying a C-terminal 6xHA epitope tag on wild-type Top2 were induced to undergo meiosis (30°C), confirming that these phenotypes are linked to *top2* and not associated with elevated temperature. Like *top2-1* mutants, *TOP2-6HA* strains exhibited substantial overlap between Hop1 and Zip1 and an appreciable number of nuclei with DAPI train tracks (**Figure S7a-c**). Interestingly, the epitope tag led to loss of Top2 from promoter-containing IGRs (**Figure S7d**), similar to *top2-1* mutants. These observations suggest that the defects in chromosome morphogenesis are related to the altered chromosomal distribution of Top2.

We asked whether the phenotypes are linked to a loss of catalytic activity of Top2. This possibility was suggested by the undetectable catalytic activity of *top2-1* mutants at 30°C (DiNardo et al, 1984) and the fact that the *TOP2-6HA* strain showed an exacerbated growth delay in the absence of *TOP1* (**Figure S7e**), indicating cells require another topoisomerase to compensate for the reduced activity of Top2-6HA. However, analysis of a catalytically inactive *top2-YF* mutant revealed no abnormalities in Hop1 removal from synapsed chromosomes (**Figure S7g**). Thus, the defects in Hop1 removal in *top2-1* mutants and *TOP2-6HA* strains are not linked to a loss in catalytic activity but may instead be the result of impaired chromosomal interactions.

Overlapping Hop1 and Zip1 signals and DAPI train tracks are indicative of a defect in proper Hop1 removal and a hallmark of mutants that fail to recruit the AAA+ ATPase Pch2 (Börner et al, 2008; San-Segundo & Roeder, 1999; Subramanian et al, 2016; Subramanian et al, 2019). The persistence of Hop1 on fully synapsed chromosomes, in turn, impairs timely DSB repair (Subramanian et al, 2016). Indeed, the number of Rad51 foci on fully synapsed chromosomes in *top2-1* mutants is significantly higher than in wild type and approaches the levels seen in *pch2Δ* mutants (**Figure 7c**).

We tested for the presence of Pch2 on *top2-1* chromosome spreads across a meiotic time course. In wild-type cells, Pch2 is strongly enriched in the nucleolus and localizes to chromosomes as soon as stretches of Zip1 form (**Figure 7d**) (San-Segundo & Roeder, 1999; Subramanian et al, 2016). In *top2-1* mutants, nucleolar binding of Pch2 occurred normally. However, Pch2 binding along chromosomes was uncoupled from SC formation and only occurred with a nearly 2-h delay. This delay led to the accumulation of fully synapsed pachytene nuclei without Pch2 staining (**Figure 7d-e**), a class of nuclei also observed in *TOP2-6HA* strains (**Figure S7f**), but not in wild type or *top1* mutants (**Figure S4d**). These data indicate that Top2 is involved in coupling Pch2 recruitment and Hop1 removal to the assembly of the SC and suggest that altered chromosomal binding of Top2 uncouples these processes.

## Discussion

Topoisomerases are essential for protecting the genome from topological stress associated with most aspects of DNA metabolism, including DNA replication, transcription, and chromosome segregation. Here, we show that topoisomerases I and II also modulate the timing of meiotic DSB formation and chromosome morphogenesis.

Our data show that similar to vegetative cells, topoisomerases are strongly enriched in IGRs during meiosis, possibly because of topological stress in these regions. Both topoisomerases exhibit preferential binding to the IGRs of highly expressed genes. In addition, spike-in analysis showed increased binding of Top2 as cells transition into meiotic prophase. It is unlikely that this increase is due to Top2 function during replication, as under our conditions, replication is complete an hour prior to the time of this analysis. An obvious candidate is the assembly of the axial element, which occurs specifically in meiotic prophase, but our analyses show that Top2 accumulates normally in *rec8* mutants, which lack axial elements (Klein et al, 1999). We observed an increase in Top2 binding at elevated temperatures, a condition that has been shown to affect meiotic chromosomes (Börner et al, 2004; Zhang et al, 2017), suggesting that Top2 binding is modulated by cellular stress. Additional stress may arise from the starvation conditions needed to induce meiosis, as nutrient depletion leads to substantial chromatin compaction in yeast (Rutledge et al, 2015). Notably, starvation-associated chromatin compaction requires the chromosome remodeler condensin (Pommier et al, 2016; Swygert et al, 2019), which frequently acts in conjunction with Top2. We therefore speculate that the combined topological stresses from changes in transcription and chromosome compaction drive topoisomerase buildup on prophase chromosomes.

Spo11, the conserved enzyme responsible for meiotic DSB formation, is structurally related to type II topoisomerases and is also most active in IGRs (Baudat & Nicolas, 1997; Pan et al, 2011). Despite these similarities, our data show spatial differences in topoisomerase binding and Spo11 activity, arguing against a single shared recruitment mechanism. These binding differences do not exclude the possibility that Spo11 also responds to DNA topology, as topological stress can propagate along the DNA fiber. Indeed, the delay in meiotic DSB initiation observed in *top1Δ* mutants may point to interplay between Spo11 and Top1. However, at this point, we cannot exclude the possibility that the *top1*-associated delay is not linked to hotspot topology but rather arises indirectly, for example from defective expression of DSB factors.

Consistent with previous results (Zhang et al, 2014), we find that inactivation of *TOP2* delays meiotic DSB turnover. The observed delays, however, are likely the combined result of multiple defects. In particular, Zhang and colleagues showed delayed DSB turnover in catalytically inactive *top2-YF* mutants, which do not detectably alter chromosome architecture. Our observations imply that altered binding of impaired Top2 can have additional effects on meiotic DSB repair. It is possible that the inactive enzyme affects repair activities, as suggested by the observation that Top2 physically interacts with the recombinase Dmc1 in *Corpinus cinereus* (Iwabata et al, 2005). Alternatively, the binding of impaired Top2 may interfere with factors involved in meiotic chromosome dynamics. An effect on chromosome architecture is supported by the fact that mutant Top2 is primarily retained at sites enriched for the chromosome axis factor Red1. This altered binding suggests that mutant Top2 is no longer efficiently recruited to sites of topological stress but may instead interact with other protein components or chromatin marks of meiotic chromosomes. In this context, it may be significant that *S. cerevisiae* Top2 shares an acidic patch at its C-terminus that functions as a chromatin-binding domain in mammalian Topo IIα (Lane et al, 2013) and that Top2 recruitment to meiotic chromosomes in *C. elegans* requires a specific chromatin-modifying enzyme (Wang et al, 2019).

Our analyses suggest that one consequence of this altered binding of Top2 is a defect in recruiting the meiotic chromosome remodeler Pch2. In wild-type meiosis, assembly of the SC coincides with removal of the HORMAD factor Hop1 from chromosomes, leading to a down-regulation of new DSB formation and an easing of meiosis-specific repair restrictions (Subramanian et al, 2016; Thacker et al, 2014). By contrast, meiotic chromosomes of *top2-1* mutants accumulate SC structures that remain decorated with Hop1, as well as “train-track” chromosomes by DAPI analysis. Both phenotypes and the concomitant delay in meiotic DSB repair are characteristic of a failure to recruit the AAA-ATPase Pch2 (Börner et al, 2008; Subramanian et al, 2016), which disassembles Hop1 from chromosomes (Chen et al, 2014). Indeed, Pch2 recruitment is notably delayed in *top2-1* mutants. These data indicate that Top2 functions upstream of Pch2 in promoting the remodeling of meiotic chromosomes during meiotic chromosome synapsis. They also indicate that even though Pch2 binding requires Zip1 deposition (San-Segundo & Roeder, 1999), Pch2 is not simply recruited by the assembly of the SC. Rather, these data point to a structural transition following SC deposition that needs to occur to promote Pch2 binding (**Figure 7f**). An appealing possibility is that this transition renders Hop1 recognizable as a substrate for Pch2, thereby leading to Pch2 recruitment. On wild-type chromosomes, this process likely occurs rapidly following SC deposition, leading to minimal overlap between Hop1-decorated axes and the SC and only transient appearance of DAPI “train-tracks”. The separation of SC deposition and structural remodeling of the SC in *top2-1* mutants is reminiscent of meiotic chromosome morphogenesis in wild-type *Caenorhabditis elegans* (Libuda et al, 2013; Pattabiraman et al, 2017). Thus, the apparent differences in synapsis between these organisms may primarily result from different pausing between SC assembly and subsequent remodeling.

Our data reveal multiple roles for topoisomerases during meiotic recombination. We speculate that this pleiotropic dependence is related to the cycles of expansion and compression of the chromatin fiber volume throughout prophase (Kleckner et al, 2004). These fluctuations are accompanied by stresses along chromosomes, including twisting and buckling, which are likely used to communicate local changes in chromosome organization (Kleckner et al, 2004). Intriguingly, the three major cycles of expansion and compression are predicted to correspond to DNA breakage, axial transitions, and untangling of chromatids. These cycles coincide well with the topoisomerase-associated phenotypes observed by others (Rose & Holm, 1993) and us and suggest that the effects of topoisomerases on the timing of meiotic prophase may result from their contribution to the volume fluctuations of meiotic chromosomes.

## Materials and Methods

### Yeast strains and growth conditions

All strains used in this study were of the SK1 background, with the exception of the SK288c spike-in strain used for SNP-ChIP analysis (Vale-Silva et al, 2019) and the *top2-1* mutant, which is congenic to SK1 (backcrossed >7x). The genotypes are listed in **Table S1**. Sequencing of the *top2-1* mutant revealed a single non-synonymous amino acid change: G829D. To induce synchronous meiosis, strains were accumulated in G1 by inoculating BYTA medium with cells at OD600 = 0.3 for 16.5 hours at 30°C (Falk et al, 2010). Cultures were washed twice with water and resuspended into SPO medium at OD600 = 1.9−2.0 at 30°C as described (Falk et al, 2010). *top2-1* cells were inoculated at OD600 = 0.8 in BYTA medium for 20 hours at room temperature. For experiments that included *top2-1* mutants, SPO cultures for all strains were washed twice with water and resuspended into SPO medium at OD600 = 1.9 at room temperature and shifted to 34°C after 1 hour.

### Chromatin immunoprecipitation

At the indicated time points, 25 ml of meiotic culture was harvested and fixed for 30 min in 1% formaldehyde. Formaldehyde was quenched by addition of 125 mM glycine and samples processed as described (Blitzblau & Hochwagen, 2013). Samples were immunoprecipitated with 2 μL of either anti-Top2 (TopoGEN, #TG2014), anti-MYC 9E11 (Abcam, #ab56), anti-HA (Roche Applied Science, 3F10, #11867423001), or anti-Red1 (kind gift of N. Hollingsworth, #16440) per IP. For SNP-ChIP experiments, previously fixed and aliquoted SK288c cells were mixed with each sample to 20% of total cell number prior to sample processing for ChIP (Vale-Silva et al, 2019). Library preparation was completed as described (Sun et al, 2015). Library quality was confirmed by Qubit HS assay kit and 2200 TapeStation. 50bp, 51bp, 75bp, or 100bp single-end sequencing was accomplished on an Illumina HiSeq 2500 or NextSeq 500 instrument. Read length and sequencing instrument did not introduce any obvious biases to the results.

### Mononucleosomal DNA preparation

At the 3-hr time point, 50 ml of meiotic culture was harvested and fixed for 30 min in 1% formaldehyde. The formaldehyde was quenched by addition of 125 mM glycine and samples processed as described (Pan et al, 2011). Library preparation and sequencing were done as outlined under the chromatin immunoprecipitation section above.

### Processing of Illumina sequence data

Sequencing reads were mapped to the SK1 genome (Yue et al, 2017) using Bowtie. Sequencing reads of 75bp or 100bp were clipped to 51bp. For paired-end sequencing, only single-end information was used. Only perfect matches across all 51bp were considered during mapping. Multiple alignments were not taken into account, which means each read only mapped to one location in the genome. Reads were extended towards 3’ ends to a final length of 200bp and probabilistically determined PCR duplications were removed in MACS-2.1.1 (https://github.com/taoliu/MACS) (Zhang et al, 2008). For data processing of mononucleosomal reads using Bowtie, bandwidth (--bw) was set to 350 for model building in MACS2, and reads were extended towards 3’ ends to a final length of 146bp. All pileups were SPMR-normalized (signal per million reads), and for ChIP-seq data, fold-enrichment of the ChIP data over the input data was calculated. Plots shown were made using two combined replicates. Mononucleosomal DNA data was combined with previously published data from (Pan et al, 2011). The 95% confidence intervals were calculated by bootstrap resampling from the data 1000 times with replacement.

### Peak Calling

To identify Top1 and Top2 protein enriched regions (peaks) for **Figure 3d**, MACS-2.1.1 (https://github.com/taoliu/MACS) (Zhang et al, 2008) was used for peak calling of the sequence data by extending reads towards 3’ ends to a final length of 200bp, removing probabilistically determined PCR duplicates and using the --broad flag to composite nearby highly enriched regions that meet the default q-value cutoff.

### mRNA preparation and sequencing

At the 3-hr time point, 1.5 ml of meiotic culture was harvested. The cells were washed in TE buffer and lysed by mechanical disruption with glass beads at 4°C. The lysate supernatant was mixed with an equivalent volume of freshly prepared 70% ethanol and purified using the RNeasy RNA isolation Kit (Qiagen). mRNA was extracted from approximately 5 μg of the total RNA samples using Sera-Mag oligo-dT beads (GE Healthcare Life Sciences). The mRNA was fragmented and used to prepare sequencing libraries according to the Illumina TruSeq Stranded mRNA sample preparation kit. Briefly, the prepared mRNA was used as a template to synthesize first-strand cDNA. Second-strand cDNA was synthesized from the first strand with the incorporation of deoxyuridine triphosphates. Finally, sequencing libraries were prepared by PCR from the cDNA samples after ligation of adapters and sequenced on an Illumina HiSeq-2500 instrument with a read length of 51 nucleotides and single-end configuration.

### RNA-seq data analysis

Single-end stranded reads were mapped to the SK1 genome assembly (Yue et al, 2017) using Tophat2 (version 2.1.1; Bowtie version 2.2.9) with first-strand library type, no novel junctions, and otherwise default options (Kim et al, 2013). Mapped reads were counted using featureCounts (from subread version 1.5.1) with default options (Liao et al, 2014). Statistical analysis was performed using a count-based workflow (Anders et al, 2013) with the edgeR Bioconductor package (version 3.12.1; (Robinson et al, 2010)). Briefly, gene counts were normalized to counts per million (cpm) reads and genes with less than 10-15 mapped reads were filtered out. Transcriptome composition bias between samples was eliminated using edgeR’s default trimmed mean of M-values (TMM) normalization. Gene length-corrected counts were calculated as transcripts per million (tpm; (Wagner et al, 2012)) and differential expression analyses were performed using the generalized linear model (GLM) quasi-likelihood F-test in edgeR. Multiplicity correction was performed with the Benjamini-Hochberg method on the p-values to control the false discovery rate.

### Mapping Spo11 oligos to SK1 genome

The Spo11-oligo raw reads were downloaded from GEO and, after combining replicates, the adaptors were clipped with fastx_clipper (fastx_toolkit/intel/0.0.14) using the following parameters: -a AGATCGGAAGAGCACACGTCTGAACTCCAGTCAC -l 15 -n - v -Q33. Reads were then trimmed using fastq_quality_trimmer with a minimum quality threshold of 20 and a minimum length of 20. The trimmed reads were mapped to the SK1 genome using BWA (bwa/intel/0.7.15), extended to 37bp, and SPMR-normalized. Peaks were identified by MACS2 using the default q-value cutoff while bypassing the shifting model. Peaks below the median signal value were discarded.

### End-labeling of Spo11-oligonucleotide complexes

End-labeling of Spo11-oligonucleotide complexes was completed as described (Thacker et al, 2014). In brief, 100 ml SPO cultures were lysed with glass beads in 10% trichloroacetic acid. Lysed cells were centrifuged and then resuspended in 1.5 ml of 2% SDS, 0.5M Tris-HCl pH 8.0, 10mM EDTA, and 2% β-mercaptoethanol. After boiling the samples, soluble protein was diluted 2X in 2% Triton X100, 30mM Tris-HCl pH 8.0, 300mM NaCl, 2mM EDTA, and 0.02% SDS. Immunoprecipitation of the Spo11-oligo complexes was performed using 2.5 μg of monoclonal mouse anti-Flag M2 antibody (Sigma). Precipitated complexes were end-labeled with 5 μCi of [α-32P]dCTP and 5 units of terminal deoxynucleotidyl transferase (Enzymatics). End-labeled complexes were run on a Bolt 4±10% bis tris plus acrylamide gel (ThermoFisher Scientific), blotted onto a PVDF membrane using an iBlot2 gel transfer device (ThermoFisher Scientific), and visualized using a Typhoon FLA 9000 (GE Healthcare).

### Chromosome spreads and tubulin immunofluorescence

Meiotic nuclear spreads were performed as described (Subramanian et al, 2016). Top2 was detected using anti-Top2 (TopoGEN, #TG2014) rabbit serum at 1:200 in blocking buffer and Alexa Fluor 488 anti-rabbit (Jackson ImmunoResearch) at 1:200. Zip1 was detected using Zip1 yC-19 goat antibody (Santa Cruz Biotechnology) at 1:200 and anti-goat Cy3 at 1:200 (Jackson ImmunoResearch). Top1-13myc was detected using anti-Myc mouse serum (Millipore 4A6) at 1:100 and FITC anti-mouse (Jackson ImmunoResearch) at 1:200. Hop1 was detected using anti-Hop1 rabbit serum (kind gift of N. Hollingsworth) at 1:200 and Alexa Fluor 488 anti-rabbit at 1:200. Pch2 was detected using anti-Pch2 rabbit antibody (kind gift of A. Shinohara) at 1:200 and Alexa Fluor 488 anti-rabbit at 1:200. Rad51 was detected using anti-Rad51 (y-180) rabbit antibody (Santa Cruz) at 1:100 and Alexa Fluor 488 anti-rabbit at 1:200. Whole-cell immunofluorescence analysis of meiotic spindles was performed as previously described (Markowitz et al, 2017), using a rat monoclonal anti-tubulin antibody (Santa Cruz Biotechnology YOL1/34) at 1:100 and FITC anti-rat (Jackson ImmunoResearch) at 1:200. Microscopy and image processing were carried out using a Deltavision Elite imaging system (Applied Precision) adapted to an Olympus IX17 microscope and analyzed using softWoRx 5.0 software.

### Southern analysis

For pulsed-field gel analysis and analysis of individual DSB hotspots by standard electrophoresis, genomic DNA was purified in agarose plugs as described (Subramanian et al, 2019). DNA was digested in-gel for analysis of DSB hotspots. Samples were melted at 65°C prior to loading. Pulse-field gel electrophoresis and Southern blotting of chromosome VIII using the *CBP2* probe was performed as described (Blitzblau et al, 2007). Analysis of the *CCT6* hotspot used a HindIII digest and a previously described probe (Thacker et al, 2014). Analysis of the *CPA2* hotspot used an XhoI digest and a probe spanning ChrX: 640,208-641,235 in the sacCer3 reference genome. Hybridization signal was detected using a Typhoon FLA 9000.

## Acknowledgements

We thank A. Amon, S. Biggins, and L. Zhang for sharing strains, N. Hollingsworth and A. Shinohara for sharing antibodies, S. Keeney for helpful discussions, and the NYU Department of Biology Sequencing Core for technical assistance and data processing.

This work was supported by the National Institutes of Health [GM123035 to AH].

## Author contributions

Conceptualization, J.H., X.S. and A.H.; Investigation, J.H., X.S., L.A.V.S., T.E.M., and A.H.; Software, J.H.; Formal Analysis, J.H. and A.H.; Resources, J.H., X.S. and A.H.; Writing – Original Draft, J.H and A.H.; Writing – Review & Editing, J.H., X.S., L.A.V.S., T.E.M., and A.H.

## Data Availability

The datasets and computer code produced in this study are available in following databases:

RNA-seq data, MNase-seq data, ChIP-seq data, and SNP-ChIP data: Gene Expression Omnibus GSE131994 ‘https://www.ncbi.nlm.nih.gov/geo/query/acc.cgi?acc=GSE131994’
Computer scripts for processing Illumina reads: Github ‘https://github.com/hochwagenlab/ChIPseq_functions/tree/master/ChIPseq_Pipeline_v3/’
Computer scripts for processing SNP-ChIP reads and calculating spike-in normalization factor: Github ‘https://github.com/hochwagenlab/ChIPseq_functions/tree/master/ChIPseq_Pipeline_hybrid_genome/’
Computer scripts for making figures: Github ‘https://github.com/hochwagenlab/topos_in_meiosis‘

