## Supplemental Material for "Topoisomerases modulate the timing of meiotic DNA breakage and chromosome morphogenesis in *Saccharomyces cerevisiae*"

**Heldrich et al., Supplemental Figure 1**

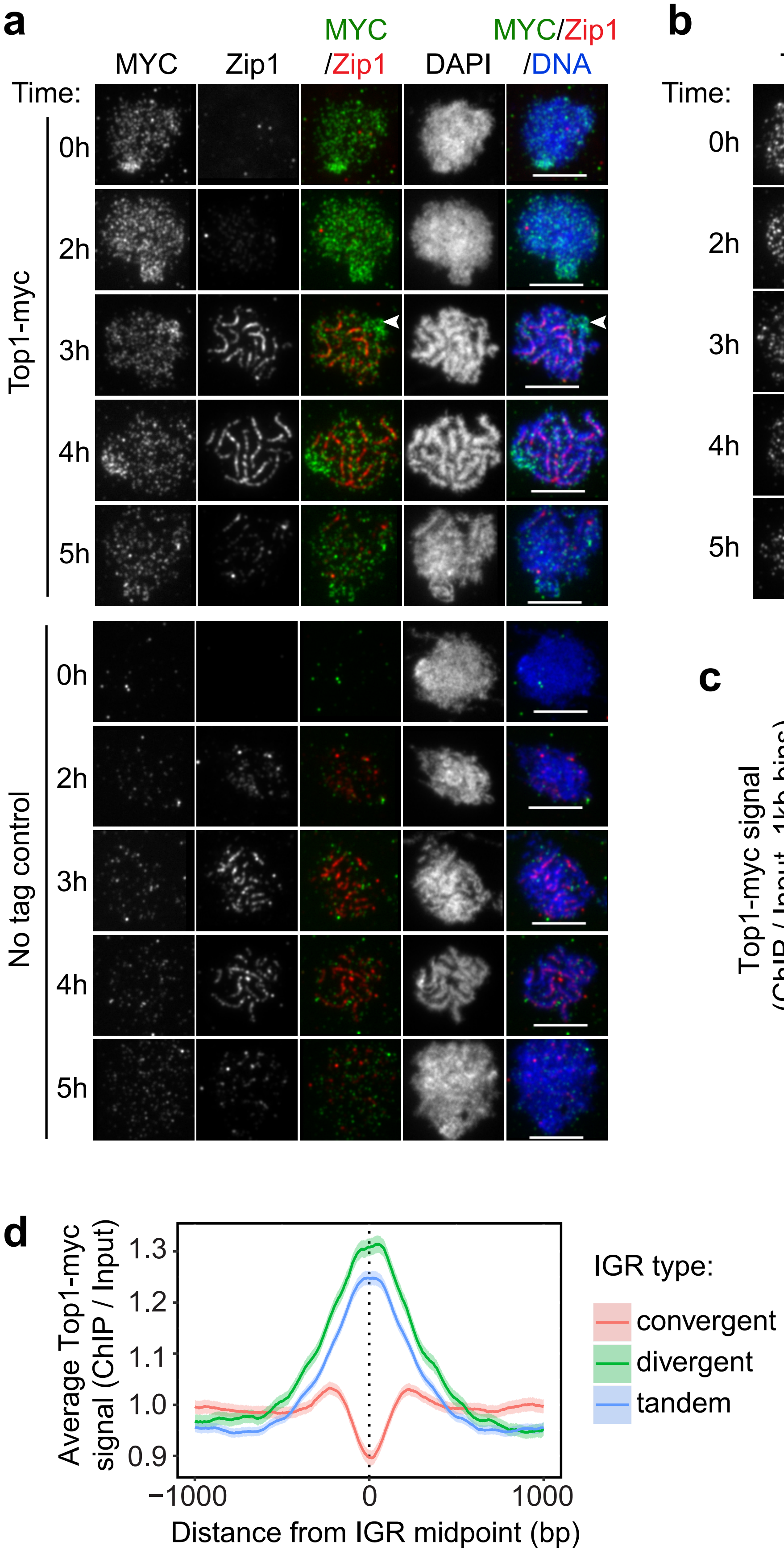

**Supplemental Figure 1. Topoisomerases are abundant on chromosomes throughout meiotic prophase.**

**(a-b)** Immunofluorescence staining of chromosome spreads throughout meiotic prophase. **(a)** Top1-13myc nuclei and wild-type no-tag control nuclei stained for the myc epitope, Zip1, and DAPI. White arrowhead marks the nucleolus. Scale bars are 5 μm. **(b)** Wild-type nuclei stained for Top2, Zip1, and DAPI. Scale bars are 5μm. **(c)** Scatterplot showing the correlation between Top1-myc and Top2 ChIP-seq signal across the genome (correlation = 0.76). **(d)** Top1-13myc localization centered at the midpoints of IGRs, parsed into divergent, tandem, and convergent regions. The 95% confidence interval is shown for average lines.

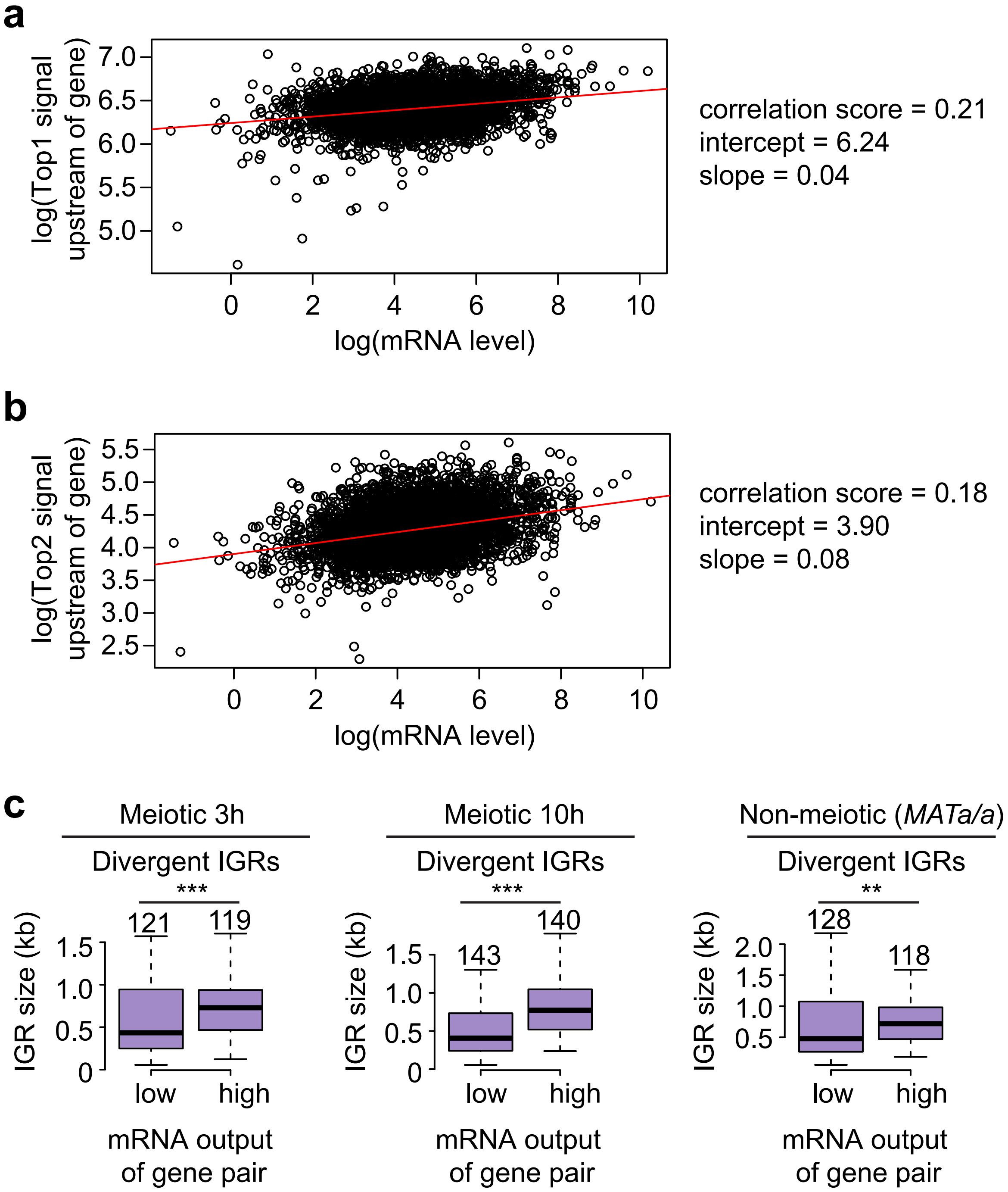
a
log(Top1 signal upstream of gene)
7.0
6.5
6.0
5.5
5.0
0
2
4
6
8
10
log(mRNA level)
correlation score = 0.21
intercept = 6.24
slope = 0.04
b
log(Top2 signal upstream of gene)
5.5
5.0
4.5
4.0
3.5
3.0
2.5
0
2
4
6
8
10
log(mRNA level)
correlation score = 0.18
intercept = 3.90
slope = 0.08
c
Meiotic 3h
Divergent IGRs
***
121
119
IGR size (kb)
1.5
1.0
0.5
0
low
high
mRNA output of gene pair
Meiotic 10h
Divergent IGRs
***
143
140
IGR size (kb)
1.5
1.0
0.5
0
low
high
mRNA output of gene pair
Non-meiotic (MATa/a)
Divergent IGRs
**
128
118
IGR size (kb)
2.0
1.5
1.0
0.5
low
high
mRNA output of gene pair

**Supplemental Figure 2. Transcription is most active in genes sharing wide divergent promoter regions during starvation.**

**(a-b)** Scatterplot comparing the relationship between mRNA level and Top1-13myc (a) and Top2 (b) signal. The values are in logarithmic scale. The red line shows the linear regression. Correlation score, intercept, and slope are noted. **(c)** Box plots showing the size distribution of divergent regions for highly and lowly transcribed gene pairs during meiosis at 3h and 10h and in non-meiotic *MATa/a* cells in sporulation medium (Cheng et al, 2018). The number of gene pairs is noted above the respective box. ***$P < 0.00001$, **$P < 0.01$, Mann-Whitney-Wilcoxon test.

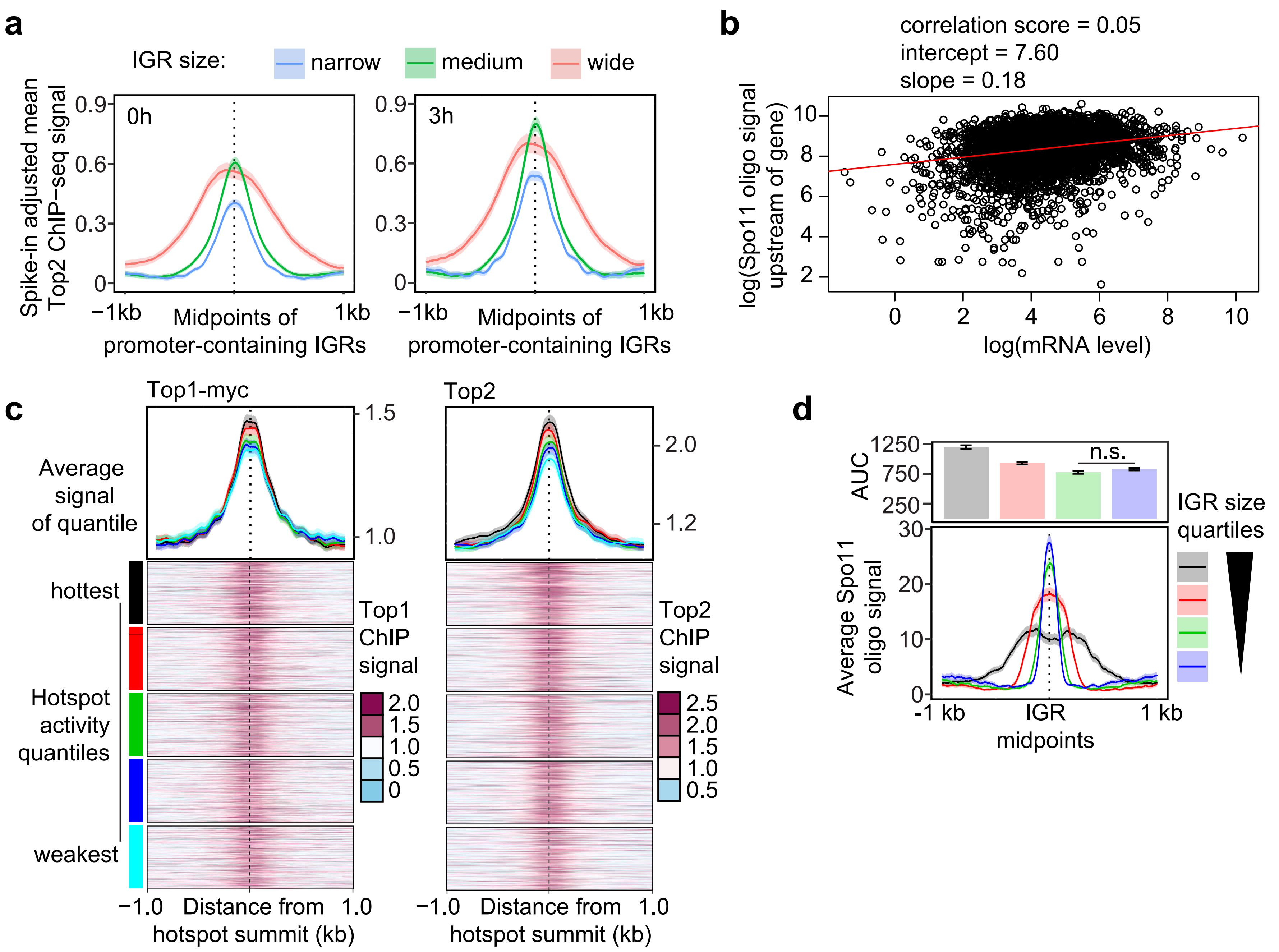
a
IGR size:
narrow
medium
wide
Spike-in adjusted mean Top2 ChIP−seq signal
0h
3h
0.9
0.6
0.3
0
−1kb
1kb
Midpoints of promoter-containing IGRs
b
correlation score = 0.05
intercept = 7.60
slope = 0.18
log(Spo11 oligo signal upstream of gene)
log(mRNA level)
c
Top1-myc
Top2
Average signal of quantile
hottest
Hotspot activity quantiles
weakest
Top1 ChIP signal
Top2 ChIP signal
−1.0
Distance from hotspot summit (kb)
1.0
d
AUC
n.s.
IGR size quartiles
Average Spo11 oligo signal
-1 kb
IGR midpoints
1 kb

**Supplemental Figure 3. Top2 abundance on chromosomes increases during meiosis.**

**(a)** Top2 binding in promoters parsed by size of the promoter-containing IGR on pre-meiotic or meiotic chromosomes using quantitative SNP-ChIP normalization. Quantile ranges are <338 bp (narrow, blue), 338-625 bp (medium, green), and >625 bp (wide, red). The 95% confidence interval for the average lines is shown. **(b)** Scatterplot comparing the relationship between mRNA level and Spo11 oligo signal. The values are in logarithmic scale. The red line shows the linear regression. Correlation score, intercept, and slope are noted. **(c)** Top1-13myc and Top2 binding centered at hotspot summits. Heat maps show all hotspots sorted by level of breakage activity. Average of each quintile is plotted above the heat maps, with the color of the line corresponding to the color beside the heat map quintile. The 95% confidence interval for the average quintile lines is shown. **(d)** Average Spo11-oligo signal in quartiles based on IGR size. Quartile ranges are <294 bp (blue), 294 - 453 bp (green), 454 - 752 bp (red), and >752 bp (black). Signals are centered at IGR midpoints and extended 1kb in each direction. The 95% confidence interval for the average lines is shown. The AUC is quantified in the bar plots above the quartile signal plot. Black bars represent standard error for each quartile. All quartiles are significantly different, except the lowest 2 quartiles ($P < 0.0001$, n.s. P-value = 0.14, Mann-Whitney-Wilcoxon test with Bonferroni correction).

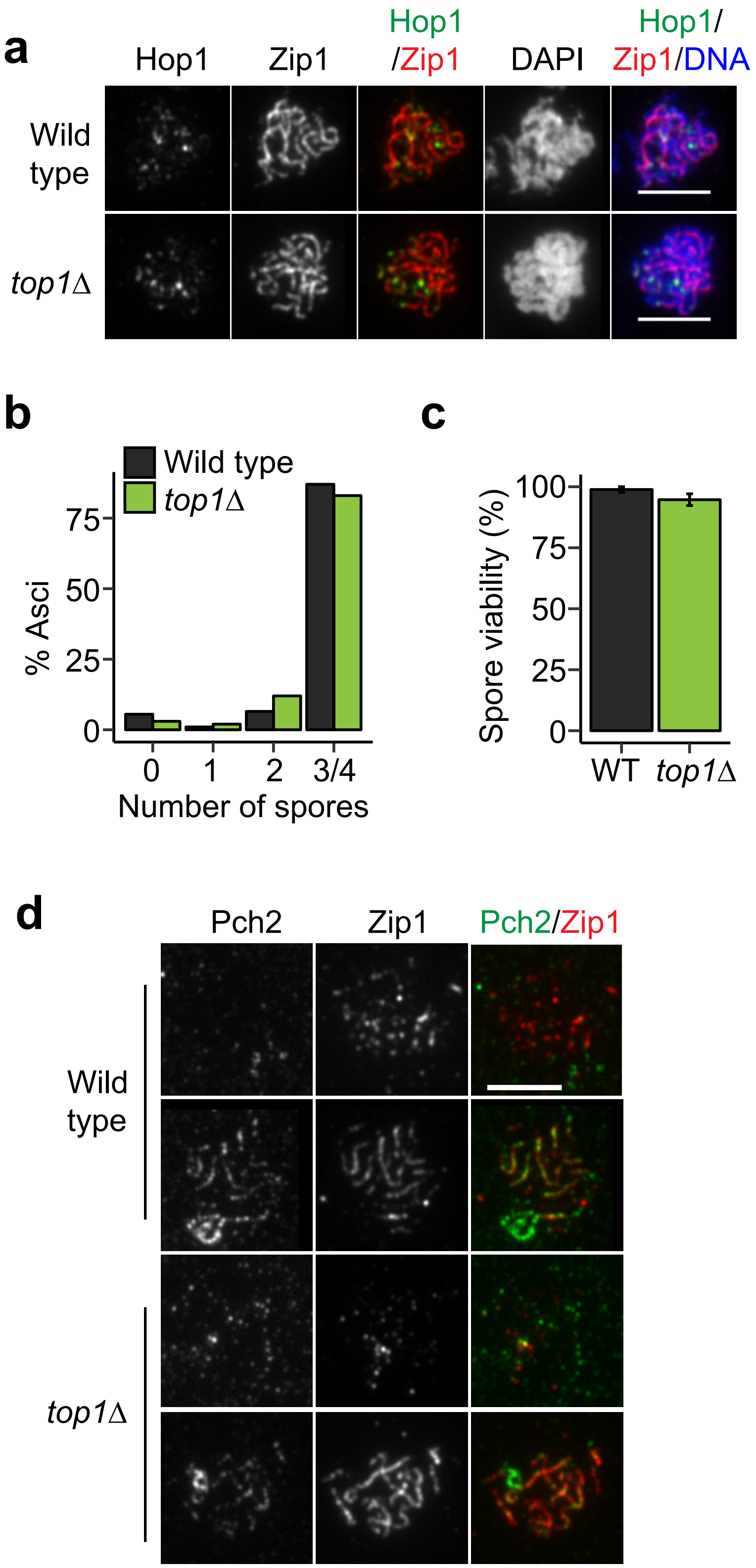

a
Hop1
Zip1
Hop1/Zip1
DAPI
Hop1/Zip1/DNA
Wild type
top1Δ
b
Wild type
top1Δ
% Asci
0
25
50
75
0
1
2
3/4
Number of spores
c
Spore viability (%)
0
25
50
75
100
WT
top1Δ
d
Pch2
Zip1
Pch2/Zip1
Wild type
top1Δ

**Supplemental Figure 4. SC morphology and sporulation are normal in the absence of Top1.**

**(a**) Immunofluorescence staining for Hop1 and Zip1 on chromosome spreads of wild type and *top1Δ* mutant during meiosis. Scale bars are 5 µm. **(b)** Sporulation efficiency for wild-type and *top1Δ* cells (n=200). **(c)** Spore viability for wild type and *top1Δ* for three experiments (n=88 per experiment). **(d)** Representative images of immunofluorescence staining for Pch2 and Zip1 on wild-type and *top1Δ* chromosome spreads 4-5 hours after meiotic induction. Scale bar is 5 µm.

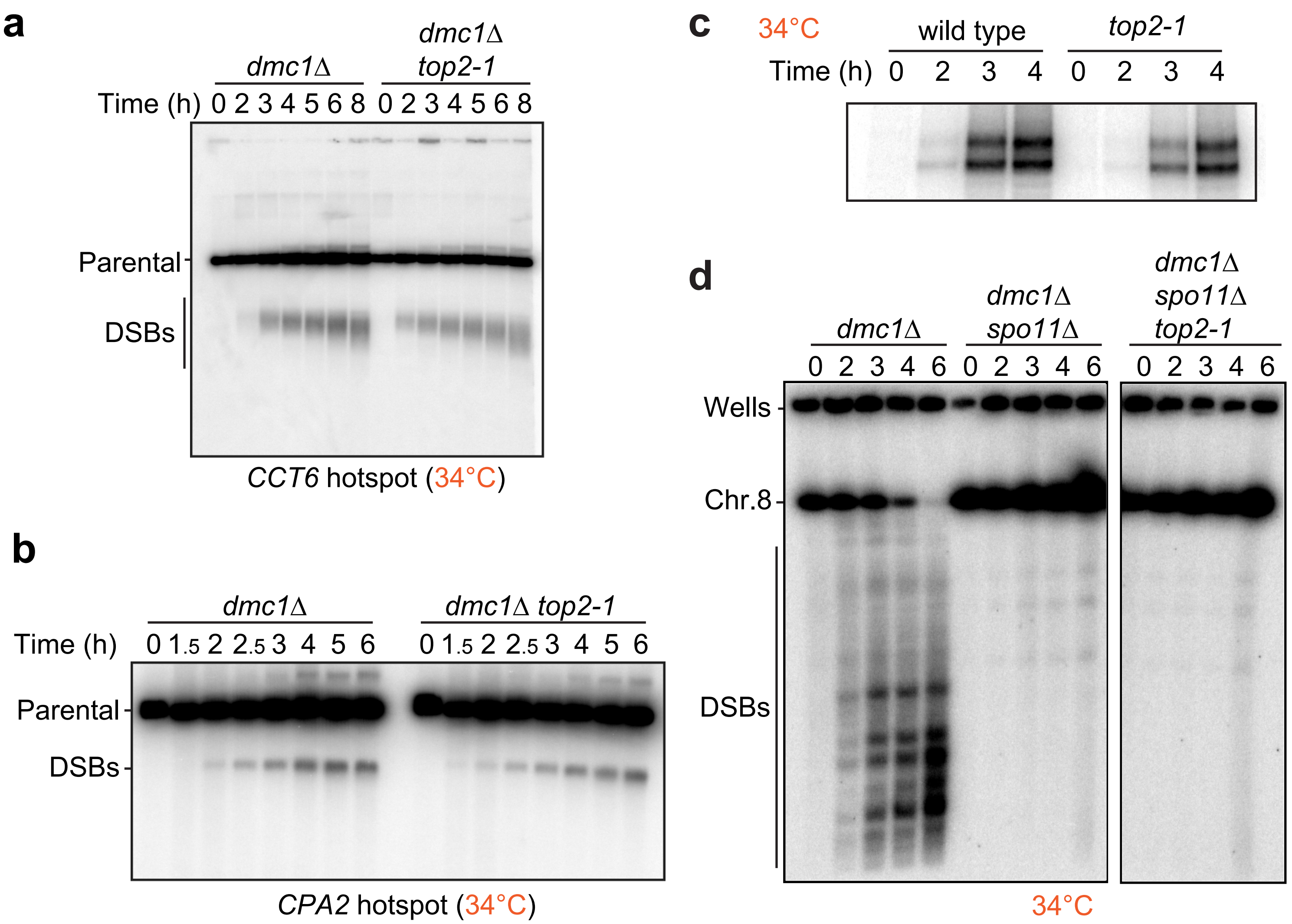
a
dmc1Δ
dmc1Δ top2-1
Time (h) 0 2 3 4 5 6 8 0 2 3 4 5 6 8
Parental
DSBs
CCT6 hotspot (34°C)
b
dmc1Δ
dmc1Δ top2-1
Time (h) 0 1.5 2 2.5 3 4 5 6 0 1.5 2 2.5 3 4 5 6
Parental
DSBs
CPA2 hotspot (34°C)
c
34°C
wild type
top2-1
Time (h) 0 2 3 4 0 2 3 4
d
dmc1Δ
dmc1Δ spo11Δ
dmc1Δ spo11Δ top2-1
0 2 3 4 6 0 2 3 4 6 0 2 3 4 6
Wells
Chr.8
DSBs
34°C

**Supplemental Figure 5. Additional analysis of DSB formation in *top2-1*.**

Southern blot analysis of the **(a)** *CCT6* and **(b)** *CPA2* hotspots throughout a meiotic time course in *dmc1Δ* and *dmc1Δ top2-1* cells. Cells were shifted to 34°C 1h after meiotic induction. **(c)** Spo11-oligo accumulation in wild-type and *top2-1* cells shifted to restrictive temperature 1 hour after meiotic induction. **(d)** Southern blot analysis of DSBs throughout a meiotic time course across chromosome VIII in *dmc1Δ*, *dmc1Δ spo11Δ*, and *dmc1Δ spo11Δ top2-1* cells. Both panels are taken from the same Southern blot image.

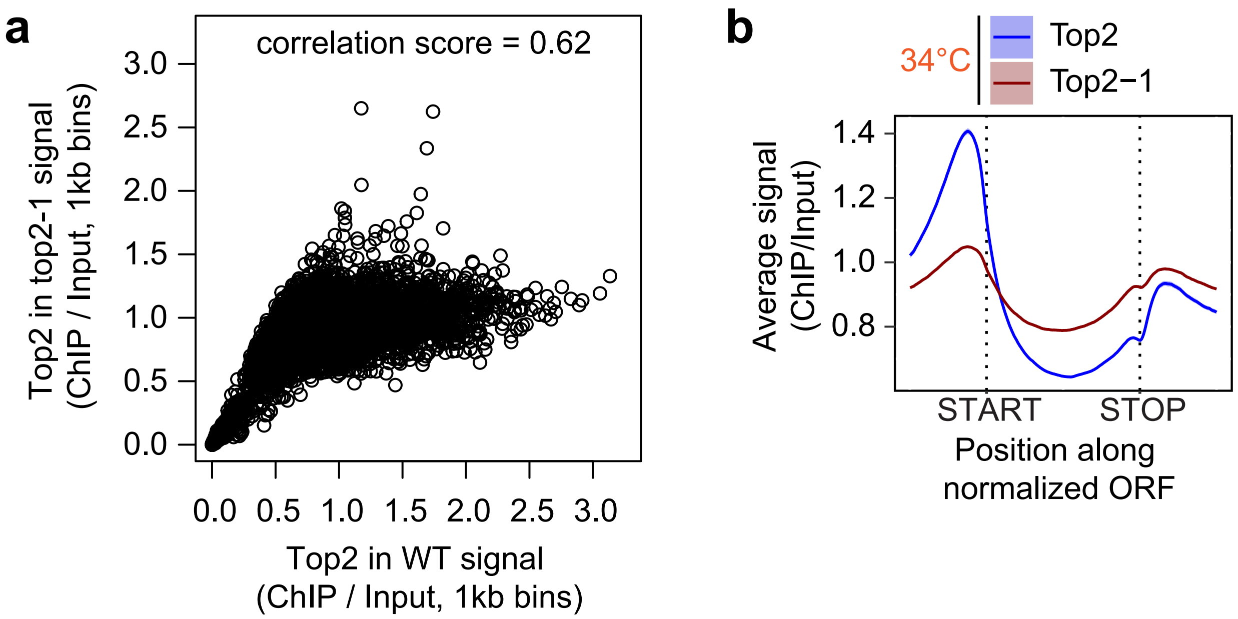
a
correlation score = 0.62
Top2 in top2-1 signal
(ChIP / Input, 1kb bins)
Top2 in WT signal
(ChIP / Input, 1kb bins)
0.0
0.5
1.0
1.5
2.0
2.5
3.0
b
34°C
Top2
Top2−1
Average signal
(ChIP/Input)
0.8
1.0
1.2
1.4
START
STOP
Position along
normalized ORF

**Supplemental Figure 6. Top2 binding is specifically lost in promoters in *top2-1*.**

**(a)** Scatterplot showing the correlation between Top2 and Top2-1 ChIP-seq signal across the genome (correlation = 0.62). **(b)** Metagene analysis of Top2 binding in wild type and the *top2-1* mutants at the restrictive temperature. The 95% confidence interval is shown for the average signals.

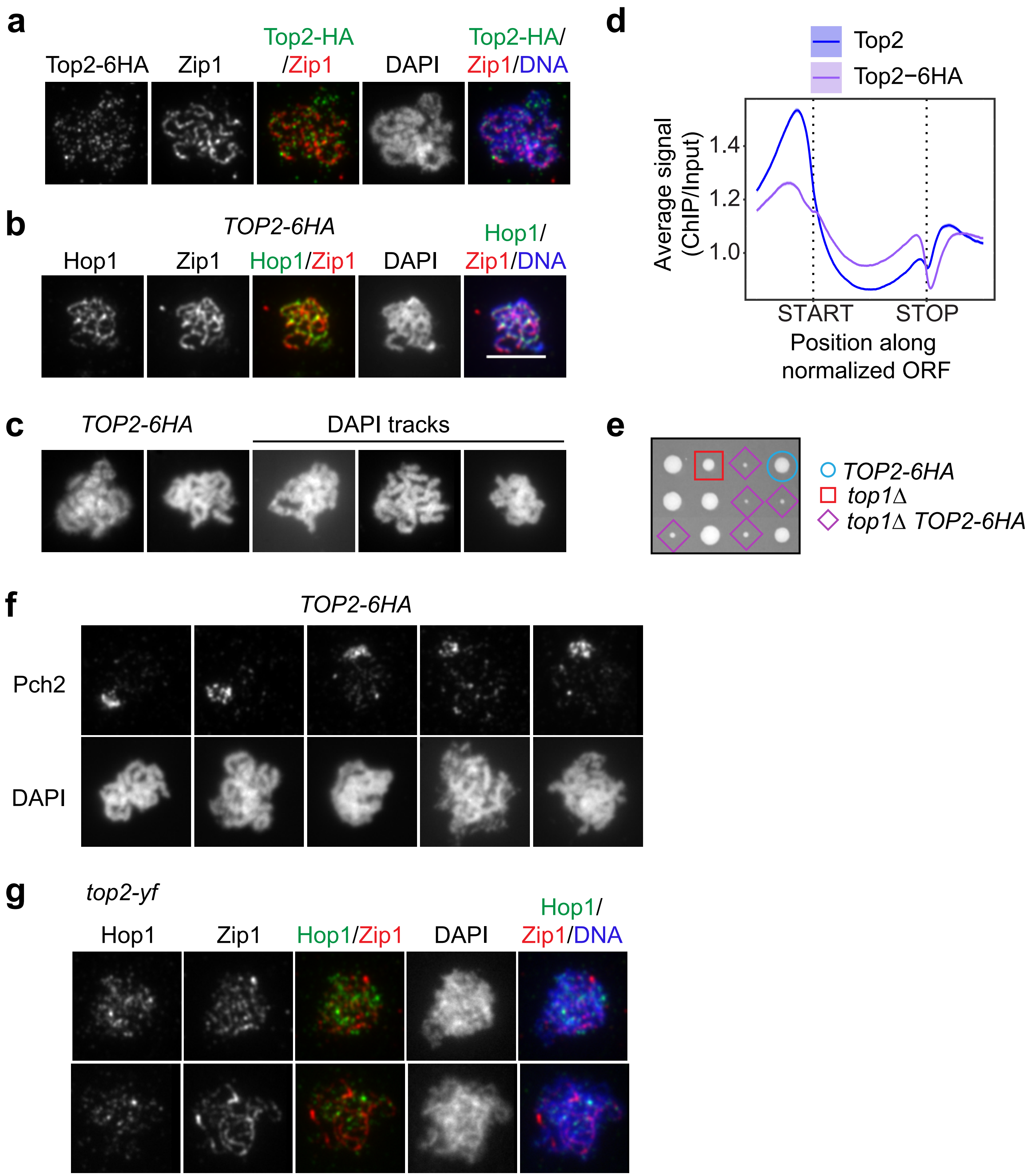
a
Top2-6HA
Zip1
Top2-HA
/Zip1
DAPI
Top2-HA/
Zip1/DNA
b
TOP2-6HA
Hop1
Zip1
Hop1/Zip1
DAPI
Hop1/
Zip1/DNA
c
TOP2-6HA
DAPI tracks
d
Top2
Top2−6HA
Average signal
(ChIP/Input)
1.4
1.2
1.0
START
STOP
Position along
normalized ORF
e
TOP2-6HA
top1Δ
top1Δ TOP2-6HA
f
TOP2-6HA
Pch2
DAPI
g
top2-yf
Hop1
Zip1
Hop1/Zip1
DAPI
Hop1/
Zip1/DNA

**Supplemental Figure 7. A C-terminal 6HA tag on Top2 interferes with meiotic chromosome morphogenesis.**

**(a)** Immunofluorescence staining for Top2 and Zip1 on chromosomes spreads of a representative *TOP2-6HA* nucleus. **(b)** Immunofluorescence staining of Hop1 and Zip1 on *TOP2-6HA* chromosomes. **(c)** Representative images of *TOP2-6HA* chromosomes as analyzed by DAPI staining. **(d)** Metagene analysis of *TOP2-6HA* at 30°C. For comparison the profile of Top2 in wild type is shown. The 95% confidence interval is shown for the average signals. **(e)** Three representative tetrads of a *TOP2-6HA* strain crossed with a *top1Δ* strain. Each horizontal quartet of colonies corresponds to a tetrad. Symbols indicate the relevant *TOP2* and *TOP1* genotypes of the respective colonies. **(f)** Example images of diminished chromosomal Pch2 signal on synapsed *TOP2-6HA* chromosomes (as determined by individualized DAPI masses). **(g)** Immunofluorescence staining for Hop1 and Zip1 on chromosome spreads of *top2-YF*. Two representative examples showing wild-type-like distribution of Zip1 and Hop1 (no overlap) are shown. Scale bars are 5 μm.

**Supplemental Table 1.** Strains used in this study

| Strain | Genotype | Background | Used in Figure Panel | Reference |
|---|---|---|---|---|
| H119 | *MATa, ho::LYS2, lys2, ura3, leu2::hisG, his4B::LEU2, arg4-Bgl II*<br>*MATalpha, ho::LYS2, lys2, ura3, leu2::hisG, his4X::LEU2 (Bam)-URA3, arg4-Nsp* | SK1 | 2b, 3b, 5d-h, 6a, 7a-b, S2a-b, S3b | (Bishop et al, 1992) |
| H7797 | *MATa, ho::LYS2, lys2, ura3, leu2::hisG, his3::hisG, trp1::hisG*<br>*MATalpha, ho::LYS2, lys2, URA3, LEU2, HIS3, TRP1* | SK1 | 1a-c, 1e, 1g, 2a, 3a, 3c-d, 4d-i, 6b-c, 7c, S1a-c, S2b, S3a, S3c, S4, S6, S7d | (Subramanian et al, 2016) |
| H118 | *MATa, ho::LYS2, lys2, leu2::hisG, his4X::LEU2-URA3, ura3, arg4-nsp, dmc1Δ::ARG4*<br>*MATalpha, ho::LYS2, lys2, leu2::hisG, his4B::LEU2, ura3, arg4-Bgl2, dmc1Δ::ARG4* | SK1 | 4a-c, 5a-c, S5a-b, S5d | (Bishop et al, 1992) |
| H8643 | *MATa, ho::LYS2, lys2, ura3::hisG, leu2::hisG, HIS4, TRP1*<br>*spo11-Y135F-HA-URA3*<br>*MATalpha, ho::LYS2, lys2, ura3, LEU2, his3::hisG, trp1::hisG,*<br>*spo11-Y135F-HA-URA3* | SK1 | 5g | |
| H10630 | *MATa, ho::LYS2, lys2, ura3, leu2::hisG, his3::hisG, arg4-Bgl II, spo11-Y135F-HA-URA3, his4B::LEU2, top2-1*<br>*MATalpha, ho::LYS2, lys2, ura3, leu2::hisG, arg4-Bgl II, spo11-Y135F-HA-URA3, his4B::LEU2, top2-1* | SK1 | 5g | |
| H9082 | *MATa, ho::LYS2, lys2, URA3, trp1::hisG, HIS3, , SPO11-6His-3FLAG-loxP-KanMX-loxP*<br>*MATalpha, ho::LYS2, lys2, ura3, TRP1, HIS3, , SPO11-6His-3FLAG-loxP-KanMX-loxP* | SK1 | S5c | (Markowitz et al, 2017) |
| H9012 | *MATalpha, ho::LYS2, lys2, ura3, leu2, his4B::LEU2, arg4-Bgl II, cyh2-z, SPO11-6His-3FLAG-loxP-KanMX-loxP, top2-1*<br>*MATa, ho::LYS2, lys2, ura3, leu2, his4X::LEU2-(Bam)-URA3, arg4, cyh2-z, SPO11-6His-3FLAG-loxP-KanMX-loxP, top2-1* | SK1 | S5c | |

| | | | |
|---|---|---|---|
| H7606 | *MATa, ho::LYS2, lys2, ura3, leu2::hisG, his4B::LEU2, arg4-Bgl II, top2-1*<br>*MATalpha, ho::LYS2, lys2, ura3, leu2::hisG, his4X::LEU2-(Bam)-URA3, arg4-Nsp, top2-1* | SK1 | 5d-h, 6, 7a-c, 7c, S6 |
| H8784 | *MATalpha, ho::LYS2, lys2, ura3, leu2::hisG, his4B::LEU2, arg4-Bgl II, dmc1Δ::ARG4, top2-1*<br>*MATa, ho::LYS2, lys2, ura3, leu2::hisG, his4X::LEU2-(Bam)-URA3, arg4-Nsp, dmc1Δ::ARG4 , top2-1* | SK1 | 5a-c, S5a-b |
| H9847 | *MATa, ho::LYS2, lys2, ura3, leu2::hisG, his3::hisG, trp1::hisG, TOP1-13Myc::TRP1*<br>*MATalpha, ho::LYS2, lys2, ura3, leu2::hisG, TRP, arg4, TOP1-13Myc::TRP1* | SK1 | 1a-b, 1d, 1f, 2a, 3d, S1a, S1c-d, S2a, S3c |
| H7652 | *MATalpha, ho::LYS2, lys2, ura3, leu2::hisG, his4X::LEU2-(Bam)-URA3, arg4-Nsp, top1Δ::kanMX6*<br>*MATa, ho::LYS2, lys2, ura3, leu2::hisG, his4B::LEU2, arg4-Bgl II, top1Δ::kanMX6* | SK1 | 4d-i, S4 |
| H7833 | *MATa, ho::LYS2, lys2, leu2::hisG, his4X::LEU2-URA3, ura3, arg4-nsp, dmc1Δ::ARG4, top1Δ::kanMX6*<br>*MATalpha, ho::LYS2, lys2, trp1, leu2::hisG, his4B::LEU2, ura3, arg4, dmc1Δ::ARG4, top1Δ::kanMX6* | SK1 | 4a-c |
| H2817 | *MATa, ho::LYS2, lys2, ura3, leu2::hisG, his4X::LEU2-(Bam)-URA3, arg4-Nsp, pch2Δ::KanMX4*<br>*MATalpha, ho::LYS2, lys2, ura3, leu2::hisG, his3::hisG, his4B::LEU2, arg4-Bgl II, pch2Δ::KanMX4* | SK1 | 7c |
| H5184 | *MATa, ho::LYS2, lys2, LEU2, ura3, trp1::hisG , his3::hisG, spo11::TRP1*<br>*MATalpha, ho::LYS2, lys2, leu2::hisG, URA3, trp1::hisG , HIS+, spo11::TRP1* | SK1 | 3a |
| H5187 | *MATa, ho::LYS2, lys2, his3::hisG, URA, leu2::hisG, TRP, rec8::HIS3MX6*<br>*MATalpha, ho::LYS2, lys2, his3::hisG, ura3, LEU2, trp1::hisG, rec8::HIS3MX6* | SK1 | 3a |
| H9173 | *MATa, ho::LYS2, lys2, ura3, leu2::hisG,his4B::LEU2, arg4-Bgl II, dmc1Δ::ARG4, spo11Δ::URA3*<br>*MATalpha, ho::LYS2, lys2, ura3, leu2::hisG, his4B::LEU2, arg4-Bgl II, dmc1Δ::ARG4, spo11Δ::URA3* | SK1 | S5d |

| | | | | |
|---|---|---|---|---|
| H9171 | *MATa, ho::LYS2, lys2, ura3, leu2::hisG, his4B::LEU2, arg4-Bgl II, dmc1Δ::ARG4, top2-1, spo11Δ::URA3*<br>*MATalpha, ho::LYS2, lys2, ura3, leu2::hisG, his4B::LEU2, arg4-Bgl II, dmc1Δ::ARG4, top2-1, spo11Δ::URA3* | SK1 | S5d | |
| H7688 | *MATalpha, ho::LYS2, lys2, ura3, leu2::hisG, his4X::LEU2-(Bam)-URA3, arg4-Nsp, TOP2-6HA::kanMX6*<br>*MATa, ho::LYS2, lys2, ura3, leu2::hisG, his4B::LEU2, arg4-Bgl II, TOP2-6HA::kanMX6* | SK1 | S7a-d, S7f | |
| H7648 | *MATalpha, ho::LYS2, lys2, ura3, leu2::hisG, his3::hisG, trp1::hisG, top1Δ::kanMX6* | SK1 | S7e | |
| H7653 | *MATa, ho::LYS2, lys2, ura3, leu2::hisG, his3::hisG, trp1::hisG, TOP2-6HA::kanMX6* | SK1 | S7e | |
| H10641 | *MATa, ho::LYS2, lys2, leu2::hisG, arg4-Nsp, ura3::Top2(Y782F)::URA3, pCLB2-TOP2::KanMX*<br>*MATalpha, ho::LYS2, lys2, leu2::hisG, his4X::LEU2-(Bam)-URA3, arg4-Nsp, ura3::Top2(Y782F)::URA3, pCLB2-TOP2::KanMX* | SK1 | S7g | |
| H8644 | *MATa, his3Δ1, LEU, LYS, ura3Δ0, RME1(ins-308a), TAO3(E1493Q), MKT1(D30G)*<br>*MATalpha, HIS3, leu2Δ0, lys2Δ0, URA3, RME1(ins-308a), TAO3(E1493Q), MKT1(D30G)* | S288C | 3a, S3a | (Vale-Silva et al, 2019) |
